# Highly contiguous genomes of *Rhodnius prolixus* and *Triatoma rubida* reveal the molecular basis of haematophagy evolution in Triatominae

**DOI:** 10.64898/2026.08.05.742999

**Authors:** Insan Habib, Carissa A. Gilliland, Hassan Tarabai, Tanisha Moons, Tyler J. Simmonds, Sheina B. Sim, Scott M. Geib, Kevin J. Vogel, Eva Novakova

**Affiliations:** Department of Parasitology, Faculty of Science, University of South Bohemia in České Budějovice, Czech Republic; Department of Entomology, The University of Georgia, Athens, GA, USA; Microbiology and Plant Pathology Department, University of California, Riverside, CA, USA; Central European Institute of Technology (CEITEC), Masaryk University, Brno, Czech Republic; USDA-ARS Tropical Pest Genetics and Molecular Biology Research Unit, Hilo, Hawaii, USA; Institute of Parasitology, Biology Centre of the Czech Academy of Sciences, Czech Republic

## Abstract

Insects of the subfamily Triatominae, commonly known as kissing bugs, are obligate blood-feeding vectors of *Trypanosoma cruzi*, the causative agent of Chagas disease. *Rhodnius prolixus* is among the most epidemiologically important vectors in Latin America, whereas *Triatoma rubida* frequently invades homes and is a potential vector in the southern United States and northern Mexico. Triatomines likely evolved from predatory reduviid assassin bugs through a transition from feeding on arthropods associated with vertebrate hosts to feeding directly on vertebrate blood. To investigate the genomic basis of this ecological and dietary shift, we generated highly contiguous, near chromosome-level genome assemblies and structural gene annotations for *R. prolixus* and *T. rubida*. The new *R. prolixus* assembly improves scaffold N50 more than 40-fold over the current reference genome, from 1.1 to 43.9 Mb, while reducing assembly gaps by several orders of magnitude. Both assemblies exceed 97% BUSCO completeness. Comparative analyses with representative hemipteran genomes revealed expansions of gene families associated with chemosensation and metabolism, including detoxification, protein degradation, and digestion, together with signatures of positive selection in genes involved in digestive and sensory functions. These assemblies represent the most contiguous and complete genomic resources available for Triatominae and provide a robust foundation for investigating vector biology, host adaptation, and the evolutionary origins of blood feeding within Reduviidae.

**Interpretive summary:** Kissing bugs are insects that are known for feeding on blood. They can spread a disease called Chagas disease because they transmit a parasite called *Trypanosoma cruzi*. To understand how kissing bugs evolved and which genes facilitate blood feeding of vertebrates, a collaboration between scientists at USDA-ARS, University of Georgia, and University of South Bohemia sequenced the genome of two kissing bugs: *Rhodnius prolixus* and *Triatoma rubida*. By comparing the genes with those of other insects in the order Hemiptera, scientists discovered that kissing bugs have more genes involved with detecting environmental chemical stimuli and metabolism as well as positive selection for genes involved with digestion and sensory-related proteins. These genome assemblies will help scientists learn more about how these insects evolved, and this research is important for understanding insect feeding biology which can be used to develop methods to control the kissing bugs and the spread of Chagas disease.

## Introduction

Kissing bugs (also known as triatomines) include approximately 150 species of blood-feeding insects belonging to the subfamily Triatominae (Hemiptera: Reduviidae), distributed almost entirely across the Americas and most diverse in the Neotropics (Schofield and Galvão 2009; Vieira, et al. 2018). Triatomines have profound medical significance as the insect vectors of *Trypanosoma cruzi*, the causative agent of Chagas disease, which continues to pose a major public health burden in the Americas. All the Chagas disease vectors are found within the subfamily Triatominae, though species vary considerably in their ability to transmit the parasite (i.e., vectorial capacity) (Stevens, et al. 2011; Echeverria and Morillo 2019).

Several species, including *Rhodnius prolixus*, are particularly effective vectors of Chagas disease due to their adaptation to domestic habitats and feeding behaviour. Owing to their large body size, tractability in laboratory settings, and medical relevance, *R. prolixus* has long been established as a model organisms in insect endocrinology (Wigglesworth 1934; Ons 2017; Lange, et al. 2022), reproductive physiology (Wigglesworth 1943), and host-microbiome interactions critical for development (Rodríguez-Ruano, et al. 2018; Brown, et al. 2020; Gilliland, et al. 2023).

In contrast to the more extensively studied *R. prolixus*, *Triatoma rubida* represents an ecologically distinct triatomine associated with arid sylvatic and peridomestic habitats in the southwestern United States and northern Mexico. In southern Arizona, *T. rubida* is closely linked to white-throated woodrat (*Neotoma albigula*) habitats, but adults disperse seasonally from late spring to early summer and frequently invade human dwellings, where they feed on humans and domestic animals (Stevens, et al. 2012; Reisenman, et al. 2014; Indacochea, et al. 2017). This behaviour is of particular medical relevance because *T. rubida* populations in the southwestern Unites States exhibit substantial *T. cruzi* infection rates, including 41.5% in Tucson, Arizona, and additional comparable reports from New Mexico and Texas (Reisenman, et al. 2010; Reisenman, et al. 2014; Indacochea, et al. 2017).

Hemipterans are an ancestrally plant-feeding group, with insectivory evolving independently in several lineages including Reduviidae. Within the Reduviidae, hematophagy (blood-feeding) is a derived trait found only in the Triatominae, though blood feeding also evolved independently in the Cimicidae. Current evidence suggests that the ancestors of Triatominae inhabited vertebrate nests or burrows, where they initially preyed on arthropods associated with these habitats before ultimately transitioning to feeding directly on vertebrate blood (Schofield 2000; Hwang and Weirauch 2012; Otálora-Luna, et al. 2015; Justi and Galvão 2017). This dramatic ecological and dietary shift required a range of corresponding evolutionary adaptations affecting morphology, host-seeking behaviour, digestion, and diuresis.

Despite their biological and medical importance, efforts to resolve the genomic basis of their transition to blood-feeding have been hindered by limited genomic resources. For nearly a decade, genomic resources were only available for a single species, *Rhodnius prolixus*. The *R. prolixus* genome, first released in 2009 and updated in 2015 (Mesquita, et al. 2015) (Accession no. GCA_000181055.3), provided a foundation for the genome architecture of a kissing bug. However, this assembly, built on Sanger and early short-read sequencing technologies, is highly fragmented, comprised of 16,537 contigs. Annotation was limited to expressed sequence tag data with lower resolution to detect rare transcripts and alternative splice variants in comparison to modern RNA-seq-based methods (Mesquita et al 2015). The fragmentation and limited annotation hinder broader macro-syntenic relationships and limits the resolution for exploring evolutionary patterns across species. Recently, long-read sequencing technologies (e.g., PacBio HiFi) paired with enriched chromosome conformation capture techniques (Hi-C) have revolutionized our ability to generate contiguous, phased, chromosome-scale assemblies even from highly heterozygous, non-model organisms (Rice and Green 2019; Rhie, et al. 2021; Feron and Waterhouse 2022).

Here, we present two high-quality triatomine genome assemblies generated from single male individuals. For *R prolixus*, we produced a substantially improved reference genome using PacBio HiFi sequencing combined with Hi-C scaffolding, resulting in the most contiguous and complete assembly currently available for the species. We also generated a *de novo* assembly of *Triatoma rubida* using PacBio HiFi sequencing and produced tissue- and stage-specific RNA-seq data that was leveraged to produce a new reference genome annotation of the *T. rubida* genome. We utilized these resources to trace the evolutionary trajectory of haematophagy within the suborder Heteroptera. Using a comparative framework that includes our assemblies along with another recently sequenced triatomine genome (*Triatoma sanguisuga*), the predatory reduviid *Rhynocoris fuscipes*, the independently evolved obligately blood feeding lineage of bed bugs (*Cimex lectularius*), and the phytophagous (*Acythrosophion pisum*) and zoophytophagous species (*Nesidiocoris tenuis* and *Halyomorpha halys*), we examined genome collinearity, gene orthology, gene family expansion and contraction, and signatures of positive selection. These analyses identify genomic changes associated with the origin of hematophagy in Triatominae and provide new insights into the evolutionary mechanisms underlying adaptation to vertebrate blood feeding.

## Materials and Methods

### Sample preparation, library construction and sequencing

High molecular weight (HMW) DNA was extracted from a portion of the thorax and abdominal tissues of a single adult *R. prolixus* and *T. rubida* male for PacBio Hifi sequencing. Samples were cryoground using a Spex GenoGrinder 2010, and DNA was extracted using the Qiagen MagAttract HMW DNA kit according to the manufacturer’s instructions. The concentration of the extracted HMW DNA was quantified using Qubit 1x dsDNA HS kit, while purity was assessed using UV-Vis spectroscopy. Fragment size-distribution was evaluated using an Agilent Femto Pulse system using Genomic DNA 165kbp kit.

Prior to library preparation, the HMW DNA was sheared to a target size of 10-15 kb using a Diagenode Megaruptor 2 (20 kbp program). Following shearing, the DNA was cleaned and concentrated using magnetic beads. PacBio HiFi libraries were then constructed using the Express Template Prep kit 2.0, including the optional post-preparation nuclease digestion step. To remove fragments shorter than 3 kb, the libraries were size selected with 40% diluted AMPure PB beads. Final library quality control was performed using the Qubit 1x dsDNA HS kit for quantification and the Agilent Femto Pulse for size verification. Sequencing was performed on a PacBio Sequel IIe system with a 30-hour movie time with 2 hours of pre-extension, utilizing the Sequel II Binding kit 2.2. HiC libraries were only generated for *R. prolixus*, using the tissue from the head and abdomen of an adult *R. prolixus.* The samples were cryoground (Spex GenoGrinder 2010) and fixed in freshly prepared TC fixation buffer, following the Arima HiC 2.0 low-input crosslinking protocol. Proximity ligation was performed using the Arima HiC 2.0 kit according to the manufacturer’s protocol.

The resulting proximity-ligated DNA was sheared using a Diagenode Biorupter Pico and size-selected to enrich for fragments between 200-600bp. The size distribution was verified on an Agilent TapeStation using High Sensitivity D5000 ScreenTapes. The Illumina compatible HiC library was then prepared using the Swift Accel NGS 2S Plus DNA library kit following the protocol outlined in the Arima HiC. Library amplification was performed using the KAPA Library Amplification Kit for 8 PCR cycles. The final libraries were quantified using the Qubit 1x dsDNA HS kit, and size-validated on the Agilent TapeStation before sequencing on an Illumina NovaSeq 6000.

Total RNA was extracted from various tissues and instars of *R. prolixus* and *T. rubida* using TriReagent (ThermoFisher) and the AllPrep RNA Kit (Qiagen), respectively, following each manufacturer’s instructions. RNA was subjected to two DNAase treatments using the Ambion TurboDNA Free kit (ThermoFisher) to remove unwanted DNA. Libraries from the extracted tissue and instar specific RNA templates (**Supplementary data: table S1**) were commercially prepared (directional mRNA library including poly-A selection) and sequenced at Novogene Europe using Illumina NovaSeq with 300 cycle chemistry and targeted output of 6 Gb of raw data per sample.

### *De novo* genome assembly, scaffolding and quality control

Vertebrate Genome Project (VGP) workflows (Rhie, et al. 2021; Larivière, et al. 2024) were utilized on Galaxy platform (Sloggett, et al. 2013) to perform assembly and quality control of *R. prolixus* and *T. rubida* genomes using the default settings in the training pipeline (Batut, et al. 2018; Lariviere, et al. 2026). Briefly, k-mer profiling of the filtered PacBio HiFi reads was performed using Meryl v 1.3 (Rhie, et al. 2020) with a k-mer length of 21. The resulting k-mer histogram was analyzed with GenomeScope2 v2.0 (Ranallo-Benavidez, et al. 2020) to infer genome properties, including estimated genome size, repeat content and heterozygosity levels. Contig assembly of *R. prolixus* and *T. rubida* HiFi reads were performed using hifiasm v0.16.1 (Cheng, et al. 2021). The primary and alternate contig assemblies were evaluated for their consensus quality (QV) using Merqury v1.3 (Rhie, et al. 2020). To remove redundant haplotypic sequences and heterozygous duplications, purge_dups v1.2.5 (Guan, et al. 2020) was utilized to identify and remove haplotigs from the primary and alternate assemblies. The purged primary assembly of *R. prolixus* was aligned to the Hi-C reads using bwa-mem2 v2.2.1 (Vasimuddin, et al. 2019) and scaffolded with SALSA v2.3 (Ghurye, et al. 2017). For *T. rubida*, only contig level assembly was retained. The scaffolded assembly for *R. prolixus* and contig level assembly for *T. rubida* were screened for bacterial content with BlobTools v1.1.1 (Laetsch and Blaxter 2017). In addition to the expected *Wolbachia*-derived sequences previously reported as integrated into the *R. prolixus* genome (Mesquita et al. 2015), contigs corresponding to symbionts of the genera *Symbiopectobacterium* and *Arsenophonus* were recovered from the *R. prolixus* and *T. rubida* metagenomic assemblies, respectively. Details of the analyses of the integrated *Wolbachia* genes and the recovered *Arsenophonus* assembly are provided in the Supplementary Methods. The mitochondrial genome for *T. rubida* and *R. prolixus* were identified from the genome assemblies using MitoHiFi (v2.0) and Mitofinder (v1.4.1), using published mitochondrial genomes (MT556664.1; NC_050328.1) as reference for assembly and annotation. Finally, sequence statistics such as NG50, N50, L50, and LG50 were generated with QUAST v5.0.2 (Mikheenko et al. 2018) and the completeness of the assemblies were assessed for completeness using BUSCO v6.0.0 (Simão, et al. 2015) against the *arthropoda_odb12* (**Supplementary methods**).

### Genome masking, RNA-seq assembly, and genome annotation

To identify repetitive elements in the *R. prolixus* and *T. rubida* genomes, de novo repeat libraries was generated for each genome using RepeatModeler v2.0.3 (Flynn, et al. 2020), followed by soft masking of complex repeats using RepeatMasker v4.2.2 (Tarailo-Graovac and Chen). RNA-seq reads from different *R. prolixus* and *T. rubida* tissues and stages were filtered using Trimmomatic (v0.39) (Bolger, et al. 2014), then aligned to their respective genome assemblies using HISAT2 v2.2.1 (Kim, et al. 2017). The resulting sorted BAM files were then used as input for Trinity v2.15.2 (Grabherr, et al. 2011) utilizing the genome-guided transcriptome assembly model to accurately reconstruct the transcriptomes of both species. Genome annotation was subsequently performed for both species using the NCBI’s Eukaryotic Genome Annotation Pipeline, EGAPx v0.4.1 (https://github.com/ncbi/egapx). For each species, a YAML configuration file was prepared containing the genome assembly, taxid, and RNA-seq evidence, and EGAPx was executed through its Nextflow workflow to generate structural annotations in GFF3 format. Finally, to carry out downstream comparative analyses, we reannotated the genomes of *N. tenuis* and *T. sanguisuga* using publicly available RNA-seq data from NCBI (BioProject: PRJNA1279500) utilizing the EGAPx pipeline, thereby ensuring consistent annotation methodology across all species included in the study (**Supplementary methods**).

### Synteny and Ortholog analysis

Genome collinearity among *R. prolixus, T. rubida* and *T. sanguisuga* genomes was investigated using SyMAP5 tool (Soderlund, et al. 2006). For each species, genome assembly FASTA files and their corresponding annotations in GFF3 format were imported into SyMAP5 and pairwise genome comparisons were performed for all three species combinations using the default alignment parameters. Synteny outputs, including *blocks.txt* and *anchors.txt*, were used for downstream visualization. Pairwise collinearity plots were generated using custom R scripts in RStudio (R version: v4.2.1), with *ggplot2* (v4.0.3), *dplyr* (v1.2.1), and *patchwork* (v1.3.2) packages (Wickham 2016; Wickham, et al. 2023; Pederson 2025) (**Supplementary methods**).

To infer orthology, protein-coding genes from genomes of eight representative heteropteran species were obtained from the NCBI genome database (**Supplementary data: table S2**). OrthoFinder v3.1.0 (Emms and Kelly 2019) was used to identify homologous gene pairs across all protein sequences. Protein FASTA files were de-isoformed first and then analysed with OrthoFinder using parameters (*-s diamond*, *-M msa*, *-A mafft*, and *-T iqtree3*). OrthoFinder outputs, including orthogroups, single-copy orthologues were used for downstream phylogenetic reconstruction, gene family evolution, and positive selection analyses. Here, an orthogroup is defined as a set of genes descended from a single ancestral gene in the last common ancestor of the species compared. Single-copy orthologues therefore contain one representative sequence per species.

The single-copy orthologous sequences clustered by OrthoFinder were used to construct a species phylogeny. First, MAFFT v7.526 (Katoh, et al. 2002) was used to align all the obtained single-copy genes. Next, TrimAl v1.5.rev1 (Capella-Gutiérrez, et al. 2009) was employed to filter out regions with unreliable homology, with *Automated1* argument employed as one of the parameters. The filtered alignments were concatenated into a supermatrix using AMAS tool v1.0 (Borowiec 2016), which grouped all the alignments into their respective species. A phylogenetic tree was constructed from this supermatrix using IQ-TREE v2.2.0 (Minh, et al. 2020). Finally, MCMCTree v4.9j (Yang and Rannala 2006) was utilized to estimate the divergence times between the species, with these estimates calibrated using fossil data obtained from the Timetree database (http://timetree.org/) (**Supplementary methods**).

### Gene family expansion and contraction

To investigate gene family evolutionacross the time-calibrated phylogeny, we used Computational Analysis of gene Family Evolution (CAFE) v5 (Mendes, et al. 2020). The orthologous group count matrix generated by OrthoFinder and the ultrametric species tree were used as input. Prior to model fitting, an error model was applied to account for non-biological variation in gene family counts arising from sequencing and annotation artefacts. We then estimated the birth-death rate (λ) under two scenarios: a single global λ shared across all branches (base model), and a model incorporating rate heterogeneity among gene families using discrete γ rate categories (γ model). The γ model was selected for final interpretation as it provided the better fit to the data. Gene families with a p-value < 0.01 were considered to have undergone significant expansion or contraction. All significantly expanded and contracted gene families were further annotated using InterProScan v5.76-107.0 (Jones, et al. 2014; Paysan-Lafosse, et al. 2023) and eggNOG-mapper v2.1.13 (Cantalapiedra, et al. 2021) using custom Python scripts (**Supplementary methods**).

### Positive selection of orthogroups

Single-copy orthologues from orthofinder were tested for evidence of positive selection. For each resulting orthogroup, protein sequences were aligned using MAFFT v7.526 (Katoh, et al. 2002), and the corresponding nucleotide sequences were codon-aligned using PAL2NAL v14 (Suyama, et al. 2006) to maintain the reading frame. Positive selection analyses were performed in Hypothesis Testing using Phylogenies (HyPhy) tool v2.5.93(MP) (Pond, et al. 2005). To test for episodic positive selection on specific branches of the phylogeny, we employed the Adaptive Branch-Site Random Effects Likelihood, aBSREL (Smith, et al. 2015), method implemented in HyPhy. Unlike standard branch-site models, aBSREL does not require a prior classification of foreground branches. Instead, it tests each branch for evidence that a subset of codon sites has evolved under positive selection (ω > 1). P-values were corrected for multiple testing using the Holm-Bonferroni method, with a significance threshold of p < 0.05.

To identify specific codons subjected to positive selection across the phylogeny, we used the Mixed Effects of Evolution, MEME (Murrell, et al. 2012), implemented in HyPhy. MEME can detect episodic positive selection by allowing the distribution of ω values to vary across all sites and among branches. Likelihood ratio tests were used to identify codons exhibiting statistically significant signatures of positive selection, with p < 0.05 used as the significance threshold. Orthogroups showing statistical significance in both aBSREL and MEME were selected for downstream functional analysis. Functional annotation of positively selected orthogroups was based on InterProScan (Jones, et al. 2014; Paysan-Lafosse, et al. 2023) and eggNOG-mapper (Cantalapiedra, et al. 2021) analyses (**Supplementary methods**).

## Results and discussion

### New genome assemblies for *R. prolixus* and *T. rubida* provide a high-resolution triatomine genomic resource

To generate a high-quality reference genome for *R. prolixus,* 2,673,715 PacBio HiFi reads were generated, yielding an average read length of 10.7 kb and a read N50 of 12.02 kb. Hi-C sequencing produced an additional 4,555,842 reads (151 bp length, 687.9 Mb total). Based on *k-*mer profiling, the genome size of *R. prolixus* was estimated to be approximately 561 Mb, exhibiting low heterozygosity (0.38%) and high sequence uniqueness (67.9%). For *T. rubida*, 1,064,146 PacBio HiFi reads with an average read length of 14.5 kb, and N50 of 15.5 kb yielded an estimated genome size of 953.9 Mb (**Supplementary data: table S3**). Contig assembly using hifiasm generated primary and secondary haplotypes of 126 and 120 contigs for *R. prolixus*, and 414 and 440 contigs for *T.* rubida, respectively, each with a GC content of 34% and 33.5%. The *R. prolixus* primary haplotype achieved an N50 of 22.28 Mb, while the secondary haplotype reached 27.97 Mb. In *T. rubida*, both haplotypes achieved N50 values of 4.11 Mb (**Supplementary data: table S4**).

Screening of the assemblies for non-host sequences recovered a near-complete genome of a *Symbiopectobacterium* symbiont from *R. prolixus* (4,034,440 bp; GC content 50.3%), as reported by Moons et al. (2026), and a draft genome of an *Arsenophonus* symbiont from *T. rubida* (3,987,882 bp; GC content 40.0%; **Supplementary Figure S1**). In addition, contigs of *Wolbachia* origin identified in *R. prolixus* were confirmed to represent horizontally transferred sequences integrated into the host genome rather than an extant symbiont (see below). Following removal of the *Symbiopectobacterium*, as well as mitochondrial scaffolds, the final *R. prolixus* assembly was resolved into 63 scaffolds covering 588.46 Mb. As no Hi-C data were generated for *T. rubida,* it was assembled at the contig level; following removal of *Arsenophonus* contigs, the final *T. rubida* assembly resolved into 394 contigs covering 934.08 Mb (**Supplementary data: table S3**). This represents the first genome assembly for this species, providing an important genomic resource for the triatominae clade and for the comparative analyses conducted in this study.

The new *R. prolixus* assembly substantially improves upon the current reference genome (GenBank GCA_000181055.3), increasing scaffold N50 more than 40-fold, from 1.09 Mb to 43.88 Mb, and extending the largest scaffold from 13.43 Mb to 68.68 Mb. The published reference spans approximately 706.82 Mb across 16,537 highly fragmented contigs and contains over 142.19 Mb of gaps (20,117.34 N bases per 100 kb), whereas our HiFi-based assembly reduces this to 1.78 ‘N’ per 100 kb (**Supplementary data: table S3**). Much of the size discrepancy between the two assemblies is therefore attributable to gap content in the older reference rather than genuine additional genomic sequence (Sarmashghi, et al. 2021). Although flow cytometry estimates place the *R. prolixus* genome between approximately 626 Mb to 753 Mb (Merle, et al. 2022), a recently deposited short-read assembly (GCA_055777405.1) exhibits a genome size highly similar to that reported here, supporting the completeness of our assembly. Consistent with this, the BUSCO analyses recovered 97.1% and 97.7% of arthropod single-copy orthologues in the *R. prolixus* and *T. rubida* assemblies, respectively, indicating that genomic content is well represented in both genomes (**Supplementary figure S2**).

A total of 252.13 Mb of repeat sequences were annotated, accounting for 42.85% of the *R. prolixus* genome. Among these, DNA transposons and retroelements were the most abundant, constituting 33.96% and 29.7% of the total repeat content, respectively. Long interspersed nuclear elements (LINEs) covered 9.55% of the total genome, followed by Tc1-IS630-Pogo family of DNA transposons at 7.5%. In *T. rubida,* repetitive sequences account for 471.2 Mb (50.45%) of the genome. Retroelements (16.93%) was the most abundant repeat sequence class with LINEs accounting for 15.43% of the *T. rubida* genome. DNA transposons and Tc1-IS630-Pogo family of DNA transposons accounted for 3.89% and 2.67% of the total genome (**Supplementary data: table S5**).

Screening of the *R. prolixus* assembly identified 129 genes or gene fragments of *Wolbachia* origin that had become integrated into the host genome through horizontal transfer. Whereas the original *R. prolixus* genome assembly reported 85 *Wolbachia*-like regions containing 27 genes, the improved assembly resolved the majority of these insertions into a single 176-kb region on scaffold 12, with the remaining loci distributed across seven additional scaffolds (**Supplementary data: table S6**). Phylogenetic analyses placed the principal insertion within the F supergroup of *Wolbachia*, clustering with symbionts associated with other *Rhodnius* species (**Supplementary Figure S3A**). Of the 129 identified *Wolbachia*-derived genes, 45 were classified as pseudogenes. Transcriptomic analyses however revealed evidence of expression for 67 integrated genes, including 29 putative pseudogenes, with most exhibiting tissue-specific transcription and the highest expression levels observed in ovaries (**Supplementary Figure S3B**). To evaluate the extent of genomic assimilation, we compared codon usage frequencies of the integrated *Wolbachia* sequences with those of *R. prolixus* and several *Wolbachia* backgrounds. Although the number of codons showing a shift depended on the reference *Wolbachia* dataset used, several codons consistently exhibited frequencies much closer to those of *R. prolixus* than to any *Wolbachia* background. In particular, CGC (Arg), GAA (Glu), and GGA (Gly) showed codon usage frequencies nearly indistinguishable from those of the host genome, while six additional codons displayed intermediate shifts toward the *R. prolixus* pattern (**Supplementary data: table S7 – S13**). Together, the widespread transcriptional activity and host-like codon usage frequencies of these loci indicate extensive assimilation of the *Wolbachia*-derived sequences into the *R. prolixus* genome.

Genome annotation, guided by RNA-seq evidence, was carried out via the EGAPx pipeline, which predicted 10,402 protein-coding genes in *T. rubida*. Whereas, in *T. sanguisuga* and *N. tenuis,* 11,880 and 10,995 protein-coding genes were predicted, respectively **(Supplementary data: table S14).** BUSCO analysis of the predicted protein set against the *arthropoda_odb12* database achieved high completeness scores, with 97.5%, 97.7%, and 94.4%, for *T. rubida*, *T. sanguisuga*, and *N. tenuis*, respectively (**Supplementary figure S2**).

### Triatomine genomes share a highly conserved synteny despite their size differences

Multi-species and pairwise collinearity analysis using SyMap was conducted across three triatomine genomes: the *R. prolixus* and *T. rubida* assemblies generated in this study, and the *T. sanguisuga* retrieved from NCBI (GCA_041753995.1). Multi-species collinearity analysis identified shared collinear blocks across all three genomes, showing genome-wide conservation across Triatominae, with syntenic regions distributed across all major *R. prolixus* scaffolds and *T. sanguisuga* contigs, and their corresponding *T. rubida* contig groups. Syntenic alignments encompassed approximately 573.8 Mb, 1.1 Gb, and 921.5 Mb of the *R. prolixus*, *T. sanguisuga*, and *T. rubida* genomes, respectively, with the remaining non-collinear regions representing lineage-specific genomic content (**Figure 1**).

**Figure 1.**
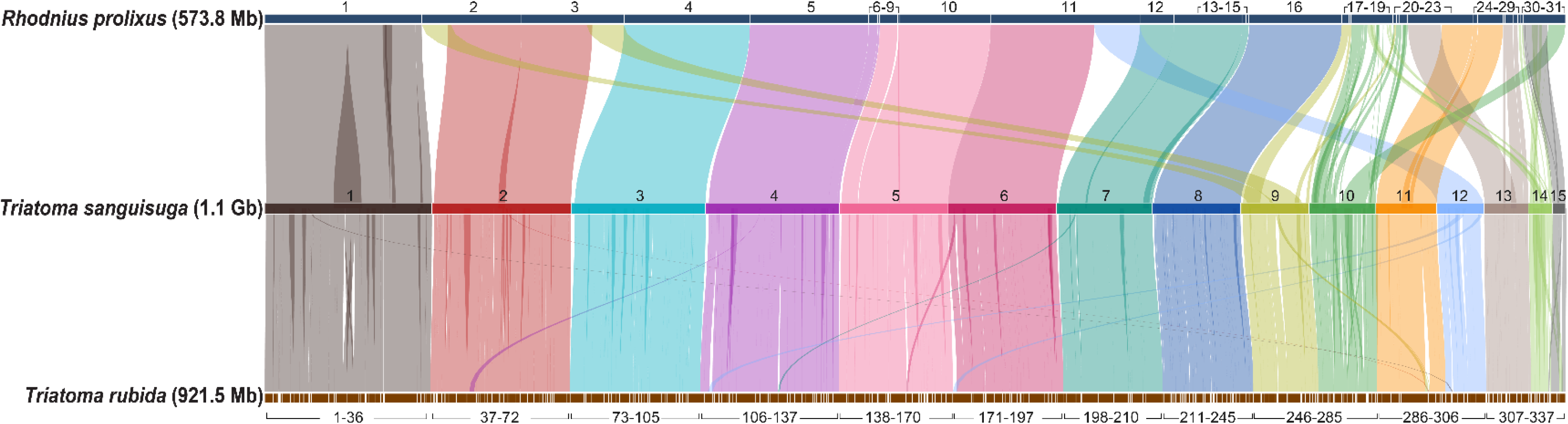
Multi-species pairwise synteny. Linear synteny plot showing genome collinearity between three triatomine species. *Rhodnius prolixus* (top), *Triatoma sanguisuga* (middle), and *Triatoma rubida* (bottom) are represented as horizontal bars, with each segment corresponding to a numbered contig or scaffold. Contig/ scaffold numbering follows order by sequence length corresponding to *T. sanguisuga*. Coloured ribbons connect collinear blocks between species, with colours also corresponding to *T. sanguisuga*. Genome size in parenthesis represents the total size of the contigs/scaffolds included in the analysis: 97.5% (573.8 Mb of 588.5 Mb) of the *R. prolixus* genome, 95.2% (1.11 Gb of 1.16 Gb) of the *T. sanguisuga* genome, and 98.7% (921.5 Mb of 934.1 Mb) of the *T. rubida* genome.

Pairwise synteny comparisons showed largely conserved 1:1 collinear relationship across major genomic regions in *R. prolixus* (572 Mb) and *T. sanguisuga* (1.1 Gb), compared to *R. prolixus* (570 Mb) and *T. rubida* (859 Mb), indicating that little large-scale structural rearrangement has occurred since their divergence. The higher degree of collinearity between *T. sanguisuga* (1.12 Gb) and *T. rubida* (922 Mb) is consistent with their phylogenetic placement as closely related lineages (**Supplementary figure S4**).

### Orthology and comparative analysis among heteropterans reveals unique genes for blood feeding adaptation

Comparative genomic analysis was conducted using OrthoFinder across three triatomines (*R. prolixus*, *T. rubida*, and *T. sanguisuga*) and five heteropterans (*A. pisum*, *H. halys*, *N. tenuis*, *C. lectularius*, *R. fuscipes*), collectively spanning haematophagous, phytophagous, and predatory dietary modes. This species selection places the triatomines within a phylogenetically informed comparative framework that enables gene family changes to be attributed to specific dietary shifts and allows the convergently evolved haematophagy of *C. lectularius* to serve as an independent reference point for identifying convergent features shared across blood-feeding lineages.

OrthoFinder assigned 94.1% (87,849 genes) of 93,391 genes into 10,758 orthogroups (hereafter OGs), with only 5.9% (5,542 genes) remaining unassigned, likely representing divergent or species-specific sequences. Of these, 4,884 (45.4%) orthogroups were shared across all eight species, reflecting the broadly conserved core gene repertoire of the Heteropterans in our study. A total of 889 (8.3%) OGs were species-specific containing 4,693 genes and likely representing potential candidate genes underlying lineage-specific adaptations, including dietary ones (**Table 1).**

**Table 1.** OrthoFinder statistics. Summary of orthology assignments across eight heteropteran species using OrthoFinder v3.1.0. The table highlights total number orthogroups, number of genes assigned to orthogroups, and single copy orthologs in all species.

| Overall Assignment | Count (%) |
| --- | --- |
| Number of species | 8 |
| Total number of genes | 93, 391 |
| Genes assigned to orthogroups | 87, 849 (94.1%) |
| Unassigned genes | 5, 542 (5.9%) |
| Total number of orthogroups | 10, 758 |
| Core and Unique Orthogroups |  |
| Orthogroups with all species present | 4, 884 |
| Single-copy orthologues | 2, 851 |
| Species-specific orthologues | 889 |
| Genes in species-specific orthogroups | 4, 693 (5.0%) |

The distribution of OGs across dietary groups is further illustrated by the UpSet plot **(Figure 2A**), which highlights several intersections, including 12 OGs shared exclusively across all blood-feeding species, the three triatomines and *C. lectularius*, but absent from *R. fuscipes* and all phytophagous species. Given that haematophagy evolved independently in the Triatominae and Cimicidae, these intersections likely represent the genes under shared selective pressure imposed by obligate blood-feeding. Similarly, 87 OGs were present across all three triatomines but absent from *R. fuscipes*. Among the 87 OGs, one OG (OG0001722) was functionally annotated as lipocalin like proteins, which contained seven copies in *R. prolixus*, two in *T. rubida*, and one in *T. sanguisuga*, suggesting conservation across the sampled triatomines together with lineage-specific differences. Lipocalins are one of the dominant and most functionally diverse protein families in triatomines, with multiple salivary subfamilies, including nitrophorins, platelet aggregation inhibitors, pallidipins, procalins, and triabins, many of which are associated with anti-haemostatic, vasodilatory, anti-platelet, or host-modulatory functions during blood-feeding (Santos, et al. 2022).

**Figure 2.**
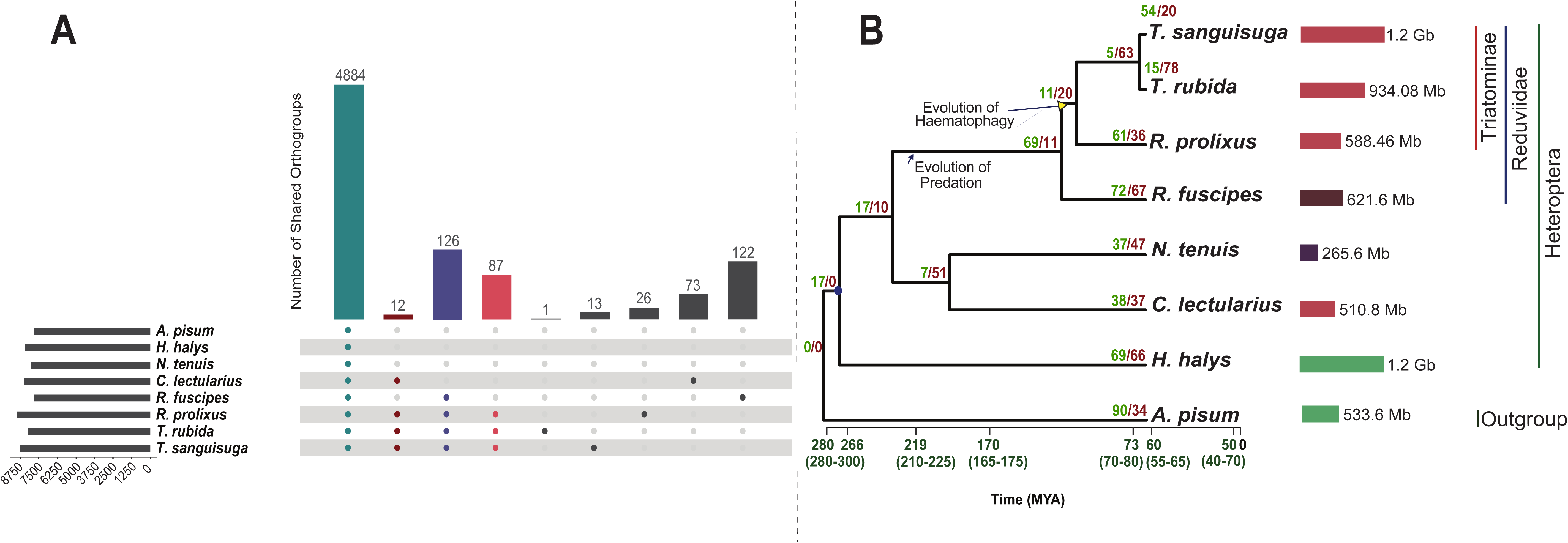
Phylogenetic relationships and orthogr oup distr ibutions acr oss eight heteropter an species. (A) UpSet plot showing the number and distribution of shared orthogroups across all eight species and blood-feeders/predators. (B) Time-calibrated phylogeny of eight heteropteran species with *Acyrthosiphon pisum* as outgroup, constructed based on previously established divergence time estimates using MCMCTree. Node bars indicate calibration intervals (in million years ago, MYA). Numbers at each node indicate expanded/contracted orthogroups (left value/right value, respectively). Colored background bars denote taxonomic transitions: plant-feeding (teal), origin of predation (light orange), and origin of blood-feeding (dark red). Genome sizes (in Mb/Gb) are indicated alongside each species name.

To determine whether the seven *R. prolixus* members of the OG were transcriptionally active, we quantified expression across our *R. prolixus* RNA-seq datasets using Salmon (**Supplementary methods**). Several members showed strong salivary-gland biased expression, with nitrophorin, triabin, procalin, and triplatin-like copies reaching high TPM values in salivary gland relative to other tissues (**Supplementary data: table S15**), which is consistent with the recent studies conducted in *R. prolixu*s, where nitrophorins, lipocalins, and triabins were enriched in salivary glands relative to *R. columbiensis* (Barbosa, et al. 2024).

We placed this OG within a broader lipocalin phylogenetic framework by extracting lipocalin-like proteins from *R. prolixus*, *T. rubida,* and *T. sanguisuga*, together with available triatomine lipocalins from NCBI. The resulting phylogeny recovered multiple known triatomine lipocalin-associated group (**Supplementary figure S5**). Notably, nitrophorin-like proteins were detected not only in *R. prolixus*, where this family was initially characterized, but also in the newly assembled *T. rubida* and *T. sanguisuga* datasets. The *R. prolixus* nitrophorins are haem-containing salivary lipocalins involved in nitric oxide transport and release and histamine binding during blood feeding. Therefore, the presence of nitrophorin-like sequences in *T. rubida* and *T. sanguisuga* suggests that this salivary lipocalin class may be more broadly distributed across Triatominae than implied by an exclusively *Rhodnius*-centered view, although functional assays would be required to confirm equivalent ligan-binding activity. In, *T. sordida*, integrative transcriptomic and proteomic analysis identified lipocalins as the most abundant family among putative secreted salivary transcripts (Praça, et al. 2022), while the *T. rubrofasciata* salivary gland transcriptome similarly found lipocalin-family proteins to be the dominant salivary component (Mizushima, et al. 2020). Similarly, in *T. rubida* sialotranscriptome analyses reported putative secreted salivary peptides including lipocalins and showed that *T. rubida* lipocalins are highly divergent from those characterized from *Hospesneotoma (Triatoma) protracta* (Ribeiro, et al. 2012). Together, the orthogroup distribution, salivary-gland expression, and phylogenetic placement support lipocalins as one of the protein families recovered in our comparative analysis and suggest that lineage-specific lipocalin diversification contributed to the vertebrate blood feeding in triatomines.

In addition to the salivary lipocalins, one metabolism-related OG (OG0009828) only present in triatomines and potentially related to their biology was annotated as a perilipin-like proteins. Perilipins are lipid-droplet associated proteins that regulate lipid storage and mobilization by influencing lipid-droplet structure and availability of stored lipolytic enzymes (Beller, et al. 2010). Because perilipins are broadly conserved regulators of lipid-droplet biology, their recovery among the triatomine-associated OGs may indicate a lineage-specific paralogue linked to lipid storage and mobilization in blood-feeding triatomines. In *R. prolixus,* substrates derived from a blood meal are converted into triacylglycerol-rich lipids in the fat body through blood-meal induced de novo lipogenesis (Saraiva, et al. 2021; Moraes, et al. 2022). Lipid mobilization is also central to post-feeding physiology in *R. prolixus*, influencing starvation resistance, reproduction, and lipid-droplet dynamics in the fat body (Braz, et al. 2023). Therefore, unlike salivary lipocalins, which are linked to host interface, perilipin-like proteins likely represent metabolic adaptations that facilitate both the storage of nutrients acquired from exceptionally large blood meals and their controlled utilization during extended periods of fasting.

The high proportion of genes assigned to OGs reflects the broad conservation of gene content across the eight heteropteran species and provides a good foundation for comparative analysis. The availability of three well-annotated triatomine genomes within this framework substantially improves the resolution of downstream comparative analyses relative to what a single triatomine reference could provide. The inclusion of the single predatory reduviid, a convergently evolved blood feeder, and phytophagous lineages in the dataset add further context, explaining whether a given OG expanded in the blood feeder or its ancestor, was gained or lost in a particular species, or changed independently across lineages.

### Gene family evolution links to molecular diversification to the origin of haematophagy

A species phylogenetic tree was constructed using 2,851single-copy orthologous sequences (1,587,675 amino acid sites), from the OrthoFinder analysis in IQ-TREE2, achieving full support (100/100 for UFB/SH-aLRT) at all nodes (**Figure 2B**). The phylogenetic analysis placed all reduviids in a single cluster, with the two *Triatoma* species forming a sister lineage to *R. prolixus*, and the entire reduviid clade sister to *C. lectularius* and *N. tenuis*. Divergence time estimation using MCMCTree with fossil calibration points from TimeTree placed the reduviid divergence from other heteropterans at approximately 219 million years ago (MYA), with the split between the predatory and blood-feeding lineages at approximately 73 MYA (**Figure 2B**). These time estimates differ from those of Hwang & Weirauch (Hwang and Weirauch 2012), who placed the origin of Reduviidae and Triatomine at approximately 178 MYA and 32 MYA, respectively, likely reflecting differences in taxon sampling as Hwang & Weirauch included a broader representation of Reduviidae and haematophagous lineages, whereas our analysis includes a single predatory reduviid representative (*R .fuscipes*).

CAFE5 identified OGs across the phylogeny, revealing contrasting patterns of expansion and contraction between haematophagous and non-haematophagous species and their ancestors. Only 191 statistically significant OGs (*p < 0.01*) were retained for downstream functional annotation and interpretation. (**Figure 2B, Supplementary data: table S16**). The predatory species, *R. fuscipes* showed the most dynamically evolving families, with 72 expansions and 67 contractions. Similarly, *H. halys*, *C. lectularius*, and *N. tenuis* showed broadly balanced profiles (69/66, 38/37, and 37/47, respectively). Within the triatomines, expansions outweighed contractions in *R. prolixus* (61/36) and *T. sanguisuga* (54/20). In contrast, *T. rubida* displayed a large reduction (15 expansions/78 contractions), whereas the convergently evolved blood feeder *C. lectularius* showed a balanced profile (38 expansions, 37 contractions) (**Figure 2B**). Significantly expanded or contracted OGs were functionally annotated using InterProScan and eggNOG-mapper and were grouped under functional categories to understand their patterns relevant to the dietary adaptation (**Figure 3; Supplementary data: table S16**), which is focused on five species, the three triatomines (*R. prolixus*, *T. rubida*, and *T. sanguisuga*), *R. fuscipes*, and *C. lectularius* to enable direct comparisons between the haematophagous and predatory lineages within the Reduviidae, and to identify features shared across independently evolved blood-feeding strategies.

**Figure 3.**
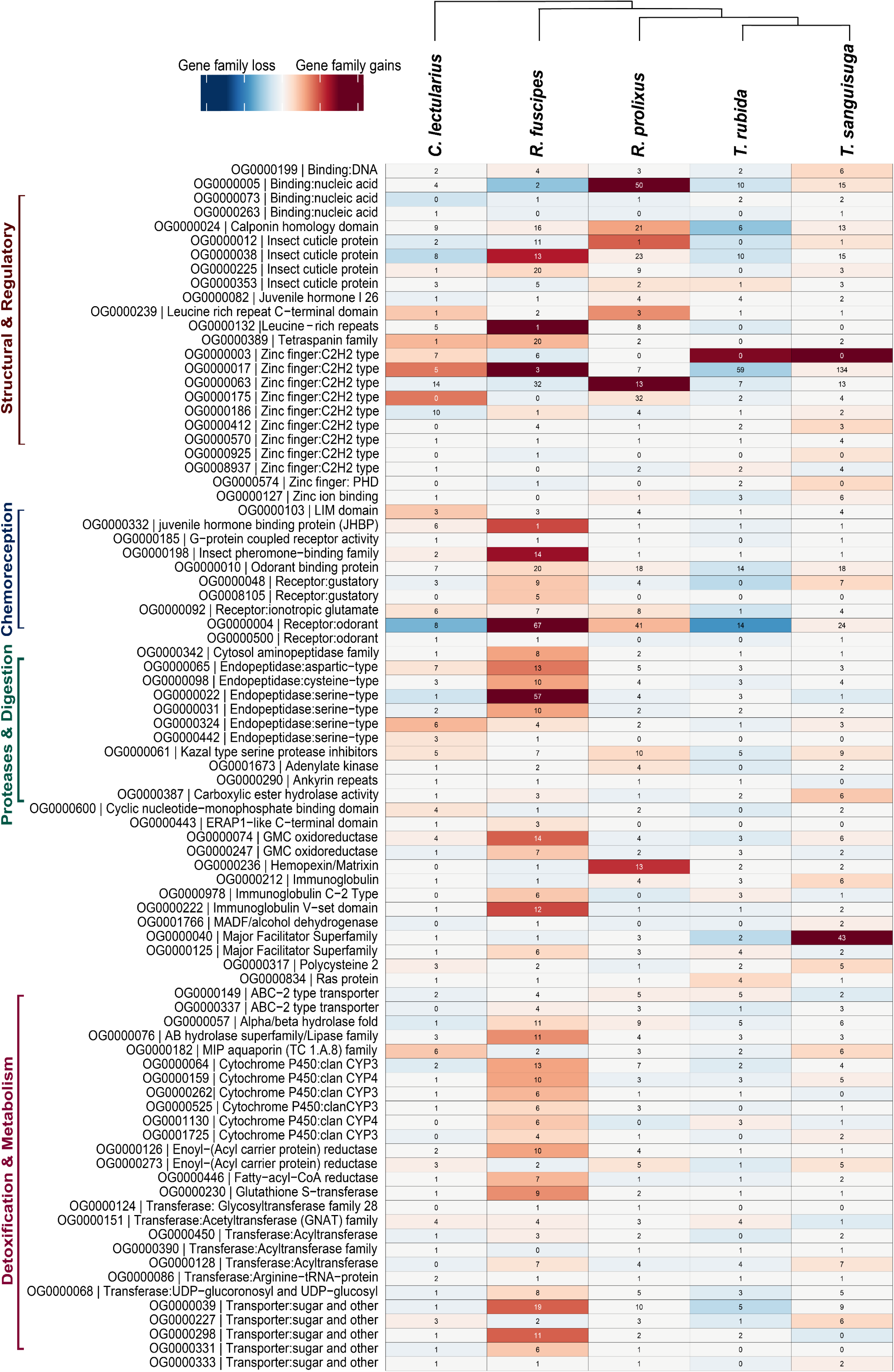
Heatmap of top expanded and contracted gene families across five heteropteran species. Gene family sizes (number of genes per orthogroup) are shown for 85 orthogroups with the most pronounced expansions or contractions across five species: *Cimex lectularius*, *Rhynocoris fuscipes*, *Rhodnius prolixus*, *Triatoma rubida*, and *Triatoma sanguisuga*. Color intensity indicates the magnitude of gene family gains (red) or losses (blue) relative to the ancestral state as inferred by CAFE. Orthogroups are grouped into four broad functional categories along the y-axis: Structural & Regulatory, Chemoreception, Proteases & Digestion, and Detoxification & Metabolism. Numeric values within cells indicate the absolute gene count per species per orthogroup.

### Contraction of chemosensory genes reflects host-seeking patterns rather than a uniform blood-feeding behaviour

Chemosensory receptor-related OGs showed varied patterns of evolution across the blood feeders (**Figure 3**). The OG (OG0000004) belonging to an odorant receptor (OR) expanded in *R. fuscipes*, *R. prolixus* and *T. sanguisuga*, but contracted in *T. rubida* and *C. lectularius,* while the OR (OG0000500) showed a moderate signal across species. The gustatory receptor (GR) OG (OG0000048) expanded in the predatory lineage but was reduced across all blood-feeding lineages, except *T. sanguisuga*. Whereas the GR (OG0008105) remained largely neutral. The ionotropic receptor (IR) OG (OG0000092) expanded in *R. fuscipes*, *R. prolixus* and *C. lectularius*, remained neutral in *T. sanguisuga*, and contracted in *T. rubida*. In addition, odorant binding proteins (OBPs) (OG0000010) showed an expansion in *R. fuscipes*, *R. prolixus* and *T. sanguisuga*, whereas it contracted in *T. rubida* and *C. lectularius*. The pheromone-binding family OG0000198 expanded only in *R. fuscipes* and *C. lectularius*, while all triatomines remained neutral.

Within the Triatominae, the chemosensory repertoire reflects their distinct host-seeking ecologies. *R. prolixus* retains the largest OR (OG0000004), likely reflecting its generalist host-seeking behaviour(Azeredo, et al. 2026). *T. sanguisuga* shows intermediate OR retention likely consistent with its broader host association (Waleckx, et al. 2014), while *T. rubida* showed more pronounced contraction, likely reflecting its more restricted ecological niche and relatively predictable host encounters in its southwestern United States (Stevens, et al. 2012; Reisenman, et al. 2014; Klotz, et al. 2016). The OR contraction in *C. lectularius* is similarly attributable to its close association with humans in a highly predictable micro-environment, where long-range host detection through diverse volatile cues is not required (Benoit, et al. 2016; Liu, et al. 2017). The reduced chemosensory related OGs in blood-feeding lineages relative to the predatory *R. fuscipes* reflect the narrower range of chemical cues relevant to an obligate blood-only diet compared to a generalist predatory lifestyle (Latorre-Estivalis, et al. 2016; Liu, et al. 2021). The broad OR expansion in *R. fuscipes* is likely consistent with the broad olfactory distinction across range of prey species (Ma, et al. 2024).

To explore the functional classes of the expanded OR copies in blood feeders, OG0000004 sequences were compared against the *Drosophila melanogaster* proteome using BLASTp via Vectorbase/VEuPathDB Several expanded copies showed homology to *D. melanogaster* odorant receptors with validated ligand in the Database of Odorant Receptors (DoOR) (Münch and Galizia 2016) to host or microbiome-associated volatile compounds (**Supplementary methods**). In *R. prolixus*, expanded copies were homologous to Or2a, for which DoOR lists 3-hydroxy-2-butanone/acetoin as the current best ligand, which is produced by cutaneous bacteria. Additional expanded copies in *R. prolixus* and *T. sanguisuga* showed homology to Or67c, Or46a, and Or49b, whose DoOR best ligands include ethyl lactate, 4-methylphenol/p-cresol and 2-methylphenol/o-cresol, respectively. Cresols are detected by olfactory sensilla or receptors in blood-feeding mosquitoes (Siju, et al. 2010; Afify and Galizia 2014), supporting their relevance as semiochemicals in haematophagous insects. However, the contracted OR repertoire in *T. rubida* included Or67c, which showed expansion in both *R. prolixus* and *T.* sanguisuga, and Or2a, expanding in *R. prolixus.* Similarly, the contracted OR copies, in *C. lectularius* (Or46a and Or49b) were expanded both in *R. prolixus* and *T. sanguisuga*. Therefore, the enrichment of OR copies homologous to receptors associated with host microbiome-derived volatile compounds specifically in the two triatomines with broader host ranges is consistent with a role for OR expansion in these lineages and has been driven for locating near vertebrate hosts, although functional assays would be required to confirm ligand specificity in triatomines.

Gustatory receptors (GRs) primarily mediate the detection of sugars, bitter compounds, CO2, and other feeding-related chemical cues that insects use to evaluate potential food sources (Rice and Green 2019; Liu, et al. 2021). GRs were reduced across both haematophagous lineages relative to the predatory and phytophagous species. Whereas, an apparent expansion of GRs was observed in *R. fuscipes*, which may be involved in determining prey suitability (Jones, et al. 2007; Ma, et al. 2024). Given the importance of CO2 perception in long-range host finding for *R. prolixus* (Otalora-Luna, et al. 2004), it is likely that CO_2_ perception is mediated by other receptors in kissing bugs.

The ionotropic receptors (IRs) are involved in detection of acids, amines, humidity, and temperature, functions extending beyond dietary chemosensation to include fundamental environmental sensing (Benton, et al. 2009; Rytz, et al. 2013; Rimal and Lee 2018). The IR copies within OG0000092 in blood feeders showed similarities with *Drosophila* Ir41a, Ir92a, and Ir76a receptors, which responds to largely amine-based ligands according to the DoOR response database. The receptor Ir76b, a co-receptor for amine detection whose orthologs in *Anopheles coluzzi* was found involved in mediating the blood-feeding behaviour (Ye, et al. 2022). Ir92a, responds to ammonia, dimethylamine, and related amines (Vulpe and Menuz 2021), and given that triatomines detects vertebrate-produced amines found in breath, urine, faeces as attractive host-associated volatiles (Taneja and Guerin 1997; Otálora-Luna and Guerin 2014; Alavez-Rosas, et al. 2024), the IR expansion in blood feeders in our study potentially reflects diversification of this amine-detection capacity, potentially serving as critical chemical cues that guide them to find and locate hosts.

OBPs were initially understood to solubilize and transport odorant molecules to olfactory receptors within chemosensory sensilla (Pelosi, et al. 2014). Subsequent studies have demonstrated that OBPs perform a wide range of functions beyond chemosensation, including driving non-sensory physiological processes in blood-feeding insects. For example, in the tsetse fly, symbiont-mediated expression of an OBP regulates host haematopoiesis and immune cell development (Benoit, et al. 2017). Within triatomines, the OBPs have additionally been implicated in blood meal processing, haeme metabolism, and xenobiotic sequestration (Santiago, et al. 2016; Oliveira, et al. 2018). The expansion and contraction patterns of OBPs (OG0000010) observed in this study likely reflect species-specific differences in host-seeking within the Triatominae and their distinct ecological niches.

### Blood feeders show a reduced detoxification capacity but an elevated transport and metabolism

Six cytochrome P450 OGs showed expansion in the predatory *R. fuscipes*, whereas obligate blood feeders exhibited contractions across several OGs (**Figure 3**). To characterize the likely functions of these expanded P450s, sequences from each OG were assigned to families using HMMER against clan and family-level p450 profiles built from an *Anopheles gambiae* P450 database (Nelson 2009) (**Supplementary methods**). Five OGs, OG0000064, OG0000262, OG0000525, OG0001130, and OG0001725, were assigned to the CYP6 family within the CYP3 clan. CYP6-family P450s are among the insect P450 lineages most frequently associated with xenobiotic metabolism and insecticide resistance. The remaining OG, OG0000159, was assigned to the CYP4 clan, whose members include enzymes involved in fatty acid metabolism, and in some cased, hydrocarbon biosynthesis (Dermauw, et al. 2020; Nauen, et al. 2022). The contraction CP450 OGs in blood-feeding lineages is consistent with reduced selection for broad xenobiotic detoxification capacity under a chemically uniform obligate-blood diet, especially when compared with the chemically diverse prey encountered by a generalist predator like *R. fuscipes* (Feyereisen 1999; Lu, et al. 2021; Nauen, et al. 2022). In parallel, reduced detoxification related genes have been reported in other dietary specialists, including the western honeybee, *Apis mellifera* (Lu, et al. 2021) and the human body louse, *Pediculus humanus* (Nauen, et al. 2022). Furthermore, transcriptomic similarities between the *C. lectularius* and *P. humanus* are directly attributed to their shared blood-feeding lifestyle (Bai, et al. 2011).

Phase II detoxification OGs showed a similar pattern to the CYP450s, glutathione S-transferase (GST, OG0000230) and UDP-glucosyl transferases (UGTs, OG0000068) expanded in *R. fuscipes* but were contracted in blood-feeding lineages. GSTs are conserved phase II enzymes that conjugate glutathione to electrophilic substrates, conferring protection against oxidative damage and facilitating excretion of toxic metabolites (Enayati, et al. 2005), whereas UGTs enhance the water-solubility of hydrophobic toxins for excretion, a vital capacity for chemically challenging diets (Bock 2016; Scott 2025). Their reduction in blood feeders is therefore consistent with the minimal selective pressure imposed by a chemically uniform blood-only diet. While blood digestion releases highly pro-oxidant free haem, blood feeders rely on targeted haem management strategies rather than GSTs. For instance, in *R. prolixus*, free haem released during blood digestion is detoxified predominantly through crystallization into non-toxic hemozoin, while additional pathways such as cysteinylglycine modification and haem-binding proteins contribute to haem/redox homeostasis (Graça-Souza, et al. 2006; Paiva-Silva, et al. 2006).

While detoxification-related OGs show contraction in blood-feeding lineages, other metabolic-related OGs expanded likely in response to physiological challenges specific to haematophagy. The expansion of the MIP aquaporin family related OG (OG0000182) in *C. lectularius* and *T. sanguisuga* represents one of the cases of convergent OG amplification in our dataset. Blood feeding imposes an acute water-balance and osmotic challenge as haematophagous insects ingest a large fluid-rich meal that must be rapidly digested after feeding. In *R. prolixus*, diuresis begins shortly after feeding and can eliminate up to half of the ingested blood-meal volume within approximately three hours (Maddrell 1964; Orchard, et al. 2021). Aquaporins are membrane water-channel proteins involved in water transport and homeostasis, and functional work in the bed bug *C. lectularius* showed that aquaporin-like genes are important for excretion, water balance, and reproduction (Tsujimoto, et al. 2017). Therefore, the independent expansion of MIP aquaporins in *C. lectularius* and *T. sanguisuga*, two independently evolved blood feeders, is consistent with adaptation for transport and homeostasis during blood-meal processing.

### Digestion-related proteins were reduced in triatomines, but anti-haemostatic inhibitors expanded

Digestion related OGs in our CAFE analysis (**Figure 3**) showed patterns consistent with the dietary shift among the species in our study. Aspartic (OG0000065), cysteine (OG0000098) and serine (OG0000022, OG0000031, OG0000324, OG0000442) endopeptidases were broadly reduced in triatomines but expanded in the generalist predator *R. fuscipes*. This predator-biased expansion is consistent with broader comparative evidence from Hemiptera showing that diet-related genes, particularly digestion related genes, are recurrently expanded in predatory lineages, whereas haematophagous species show a lower fraction of expanded diet-related OGs that predatory species (Ma, et al. 2025). However, *C. lectularius* showed isolated expansion of only two of the serine endopeptidases OGs (OG0000324 and OG0000442). The kazal-type serine protease inhibitors (OG0000061) showed a pattern of expansion in blood feeders, particularly *R. prolixus*, *T. sanguisuga*, and *C. lectularius*, while reduced in *T. rubida*.

These patterns are consistent with the specialized digestive physiology of blood-feeding Hemiptera. Most insects rely primarily on midgut serine proteases, such as trypsins and chymotrypsins for protein digestion whereas triatomines utilize an acidic cathepsin-like proteases, including cathepsin D-like aspartic proteases and C1 family cysteine proteases (Henriques, et al. 2017; Ouali and Bousbata 2024). Therefore, the lower copy numbers of several digestive protease OGs in triatomines should not be interpreted as reduced digestive capacity, but rather as a shift toward a more specialised proteolytic system for haemoglobin digestion. This contrasts with the broader protease expansion in *R. fuscipes*, which likely reflects the more diverse protein substrates encountered by a generalist predator (Henriques, et al. 2017). A similar distinction was observed by Ma et al. (Ma, et al. 2025), who found recurrent expansion and fast evolution of digestion-related genes such as trypsin and carboxypeptidase in predatory Hemipterans, supporting the idea that predatory feeding is associated with broader digestive gene recruitment. However, for triatomines, this transition may have been facilitated by the evolutionary loss of the peritrophic membrane in ancestral hemipterans (Gutiérrez-Cabrera, et al. 2016; Henriques, et al. 2017).

The expansion of Kazal-type serine protease inhibitors in blood-feeding lineages reflects the unique physiological demands imposed by feeding on a living host. Blood feeding requires insects to counteract host haemostasis, including coagulation, platelet aggregation, and vasoconstriction, so salivary and gut-associated protease inhibitors are frequently recruited as anti-haemostatic factors in haematophagous arthropods. In *R. prolixus*, rhodniin is a Kazal-type serine protease inhibitor that binds thrombin to maintain blood fluidity during feeding and early stages of blood-meal processing (Friedrich, et al. 1993; van de Locht, et al. 1995; Rimphanitchayakit and Tassanakajon 2010; Ribeiro, et al. 2014). However, Kazal inhibitors are not exclusively anticoagulants. Other inhibitors, like RpTI (*Rhodnius prolixus* trypsin inhibitor), a trypsin inhibitor modulated by *Trypanosoma cruzi*, do not act as anticoagulants but instead regulate the anterior midgut microbiota, preventing bacterial overgrowth to facilitate parasite survival (Soares, et al. 2015).

### Structural and regulatory proteins vary by species rather than by diet

Several structural and regulatory protein families showed heterogeneous patterns of expansion and contraction, suggesting lineage-specific evolution rather than a single feeding-mode associated signal. Several insect cuticle protein OGs, Zinc finger:C2H2, leucine-rich repeats, tetraspanins, juvenile hormone-binding proteins, and LIM/calponin homology domain proteins OGs varied among species, but most did not exhibit consistent expansion across all blood-feeding lineages in our dataset.

However, they remain biologically important. For example, cuticle proteins, which showed expansions particularly in *R. prolixus*, *T. sanguisuga*, and *R. fuscipes*, are major structural components of the arthropod exoskeleton, and blood-feeding insects experience strong mechanical demands during engorgement and moulting (Willis 2010). In *R. prolixus* and *Triatoma infestans*, blood feeding is associated with rapid plasticization of the abdominal cuticle, allowing the body wall to expand during engorgement (Ianowski, et al. 1998). Similarly, in *C. lectularius*, blood feeding triggers cuticle renewal during moulting (Flaven-Pouchon, et al. 2024).

Similar patterns were seen in insects for the calponin-homology domain OG in our dataset and no uniform expansion or reduced pattern were seen. Because calponin-homology and LIM domains are conserved actin-associated cytoskeletal proteins involved in cellular architecture in invertebrates (Galkin, et al. 2006), this pattern may reflect species-specific differences in tissue architecture rather than diet alone. Finally, the expansion of the juvenile hormone binding protein OG (OG0000332), among blood feeders, was only observed in *C. lectularius*, where juvenile hormone signalling is known to regulate vitellogenesis and oogenesis in bed bugs (Gujar and Palli 2016); however, this pattern was not shared by the triatomines. Overall, these structural and regulatory related OGs examined here, are likely correlated with species-specific developmental, cytoskeletal, and regulatory diversification, rather than with blood-feeding adaptation.

### Positive selection is largely lineage-specific rather than shared across blood feeders

Positive selection analysis using HyPhy aBSREL and MEME identified a high-confidence set of 82 OGs showing evidence of episodic positive selection across blood-feeding lineage and their nodes (**Supplementary data: table 17**). Prior to mapping these candidates onto the species phylogeny (**Figure 4**), OGs were functionally annotated using Interproscan and eggNOG-mapper, and manually assigned to six functional categories: structural and regulatory proteins, metabolism and detoxification, proteases and digestion, sensory and signalling, vesicle trafficking, and unclassified proteins based on eggNOG-mapper’s COG identifier, with Cluster of Orthogroups database (https://www.ncbi.nlm.nih.gov/research/cog<u>#</u>).

**Figure 4.**
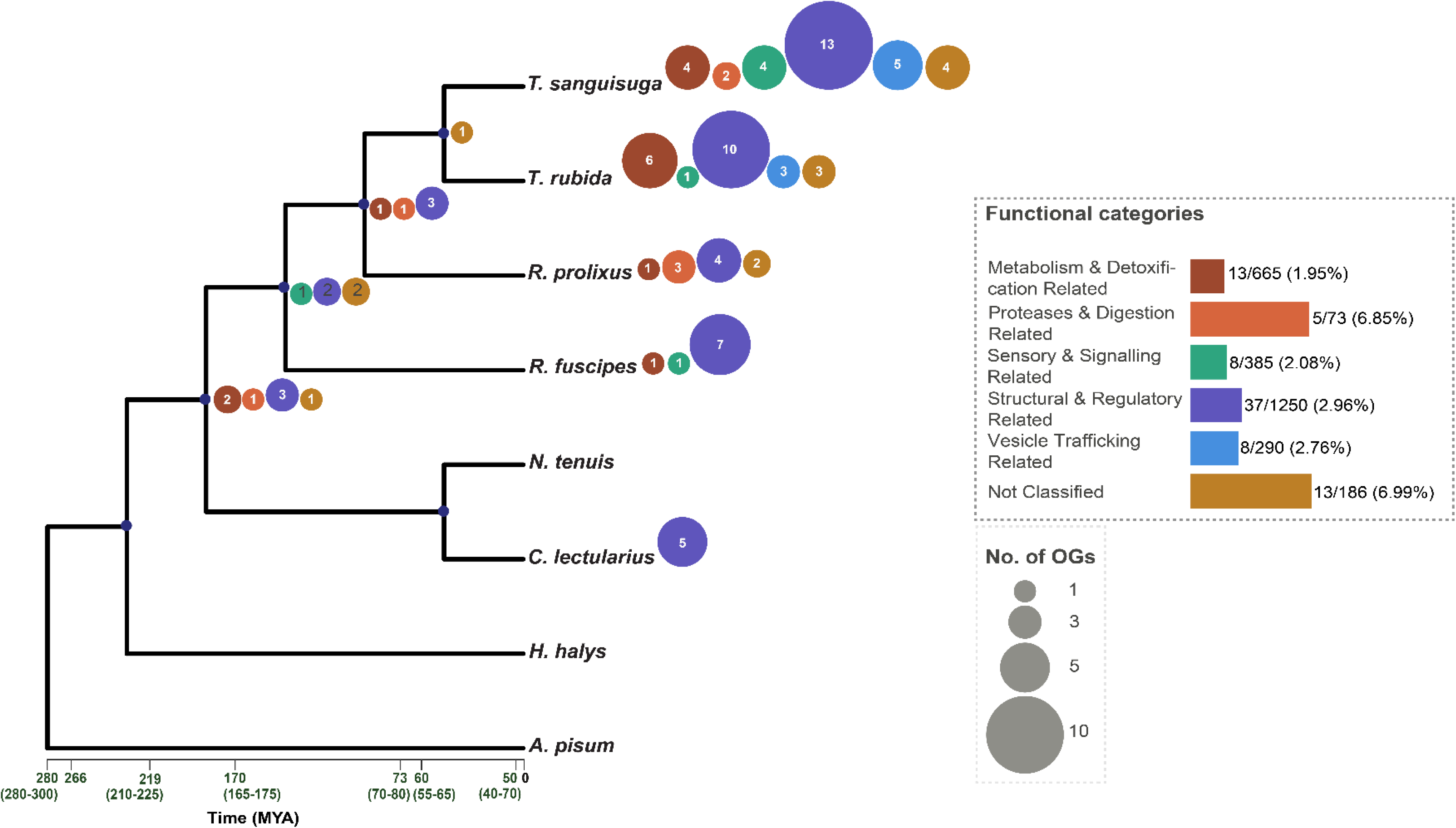
Lineage-specific distribution of orthogroups under positive selection across blood-feeding and predatory lineages. Positively selected orthogroups were identified from 2851 single-copy orthogroups using HyPhy (aBSREL + MEME) analyses. Functional categories were assigned based on Cluster of orthogroups database from eggnog-mapper output. Coloured circles indicate positively selected orthogroups on each terminal lineage or their ancestral node, with colours corresponding to the functional category legend shown on the right along with a bar chart. Circle size and the number inside each circle represent the number of positively selected orthogroup in that category. The bar chart summarizes the overall distribution of positively selected orthogroups; numbers beside the bars indicate positively selected orthogroups / total tested orthogroups assigned to that category.

Orthogroups related to structural and regulatory proteins comprised the dominant category under positive selection across all focal lineages, with the strongest signals in *T. sanguisuga* and *T. rubida*, followed by *R. prolixus* and *R. fuscipes*. In contrast, *C. lectularius* showed a more restricted positive selection signal in our dataset, confined primarily to the structural and regulatory category. Whereas proteases and digestion-related OGs showed positive selection across multiple triatomine lineages and at the ancestral node of *Triatoma* and *Rhodnius.* The sensory and signalling OGs showed modest but consistent signals in both the *Triatoma* species and the predatory bug along with the ancestral node of predatory and blood-feeding lineages. Finally, vesicle trafficking OGs showed positive selected in *T. sanguisuga* and *T. rubida* specifically. Overall, this distribution of positively selected OGs suggests a largely lineage-specific rather than a single shared adaption among blood feeders. Therefore, these results do not point to one major OG universally associated with the evolution of blood-feeding in our dataset, but rather that different reduviid and cimicid lineages likely evolved distinct molecular solutions to the challenges posed by obligate blood feeding.

Nevertheless, several positively selected orthogroups are functionally relevant to blood-feeding or feeding-associated physiology. For example, among the proteases and digestion related groups, a trypsin-like serine protease (OG0004668 & OG0004718) showed positive selection in *R. prolixus*, at the ancestral node of *Triatoma* and *Rhodnius*, and the ancestral node joining cimicid and reduviid lineages represented in this dataset (**Supplementary data: table 17**). Proteases are central to the biology of the haematophagous arthropods, where they contribute to blood meal processing (Isoe, et al. 2009; Santiago, et al. 2017). However, in triatomines, haemoglobin digestion relies strongly on aspartic and cysteine peptidases rather than trypsin/chymotrypsin peptidases as seen in many arthropods (Ouali and Bousbata 2024), a trypsin-like selection in this dataset might reflect a potentially important feeding-associated aspect. Similarly, a Zinc carboxypeptidase orthogroup (OG0005505; **Supplementary data: table 17**) was positively selected in *R. prolixus* and *T. sanguisuga*. Carboxypeptidases participate in the terminal steps of protein digestion by releasing amino acids from peptide substrates. In *R. prolixus*, haemoglobin digestion is initiated mainly by aspartic proteases and cathepsin B-like cysteine peptidases and is subsequently continued by aminopeptidases and carboxypeptidases (Ouali and Bousbata 2024). The positive selection of this OG in two blood-feeding triatomines therefore provides a plausible digestive candidate, likely specific to triatomines.

Finally, the TRP-channel orthogroup (OG00004581; **Supplementary data: table 17**) under sensory and signalling related proteins, positively selected at the ancestral node of triatomines and *R.* fuscipes is notable because TRP channels are an important sensory receptor in insects that mediate thermal and chemical sensing in insects (Montell 2005). Heat is a major host-associated cue for kissing bugs and other blood-sucking insects (Fresquet and Lazzari 2011; Lazzari 2019), and recent work has shown that TRPA5 in *R. prolixus* functions as a highly thermosensitive receptor (Liénard, et al. 2024). Thus, the selection of TRP ion associated channels among the two reduviids is likely consistent with adaptation for host or prey detection, rather than a selection specifically for blood. Taken together, the positive selection results in our dataset do not identify a single molecular route to haematophagy. Instead, they reveal a mixed pattern in which most signals are lineage-specific, while a smaller subset of digestion and sensory-related proteins correlates with feeding, digestion and sensory adaptations

## Conclusion

The transition from predation to obligate blood feeding represents one of the remarkable ecological shifts in insect evolution. By generating the most contiguous and complete genomic resources currently available for Triatominae and placing them within a comparative framework spanning diverse heteropteran feeding strategies, we present an unprecedented foundation for investigating the genomic basis of haematophagy evolution in kissing bugs. Our analyses reveal that the evolution of blood feeding was not driven by a single defining trend but instead involved a mosaic of adaptations affecting multiple biological systems. Expansions of chemosensory, digestive, detoxification, and metabolic gene families, together with lineage-specific signatures of positive selection, indicate that different triatomine lineages have followed partially distinct evolutionary routes while converging on a common blood-feeding lifestyle. The identification of triatomine-associated candidates, including salivary lipocalins and perilipin-like proteins, further highlights the interplay between host exploitation, nutrient processing, and long-term energy management that characterizes hematophagy. Beyond advancing our understanding of blood-feeding evolution, the resources generated in this study strengthen a foundation for future investigations of vector competence, host preference, and physiological adaptation in kissing bugs. As major vectors of *Trypanosoma cruzi*, triatomines remain of considerable medical importance, and these genomic resources will facilitate efforts to link genetic variation with traits relevant to disease transmission and control. More broadly, they provide a framework for understanding how predatory reduviid ancestors evolved into specialized vertebrate blood feeders and for dissecting the genetic mechanisms underlying one of the most consequential dietary transitions in insect evolution.

## Acknowledgements

This work was supported by the Czech Science Foundation (grant number 21-10185M to EN) and by the United States Department of Agriculture, Agricultural Research Service as part of the USDA-ARS Ag100Pest Initiative with funding from USDA-ARS project 2040-30400-003-000-D and USDA-ARS SCINet projects 0201-88888-003-000D and 0201-88888-002-000D. Mention of trade names or commercial products in this publication is solely for the purpose of providing specific information and does not imply recommendation or endorsement by the U.S. Department of Agriculture. USDA is an equal opportunity provider and employer.

## Author Contributions

E.N. and K.J.V. conceived and conceptualized the study. I.H. and H.T. performed the bioinformatic and comparative genomic analyses, interpreted the results, and drafted the original manuscript. T.J.S., S.B.S., and S.M.G. generated the PacBio HiFi and Hi-C sequencing data and produced the initial genome assemblies. C.A.G. and T.M. contributed to data analysis and interpretation. All authors contributed to manuscript writing, reviewed, edited, and approved its final version.

## Data Availability

All supplementary data and analysis outputs and additional genome annotations supporting this study are deposited in the Zenodo repository at https://doi.org/10.5281/zenodo.21286474. Genome assembly for *T. rubida* are deposited at NCBI under BioProject PRJNA1473261 and accession number JCAHRQ000000000. All the custom scripts related to this study are submitted in https://github.com/Thenameisinsan/Triatominae-comparative-genomics.

