## Supplementary methods for "Highly contiguous genomes of *Rhodnius prolixus* and *Triatoma rubida* reveal the molecular basis of haematophagy evolution in Triatominae"

**S1. Overview of computational workflow**

Genome assembly, annotation, orthology inference, gene-family evolution, and positive-selection analyses were performed using multi-step bioinformatics workflow. PacBio HiFi reads were used for de novo contig assembly of *Rhodnius prolixus* and *Triatoma rubida*. Hi-C data were available for *R. prolixus* and were used for scaffolding, whereas *T. rubida* was retained as a contig-level assembly. Repetitive elements were identified de novo and soft-masked before RNA-seq guided transcriptome assembly and structural genome annotation. For downstream comparative genomics, predicted protein sets of *Triatoma sanguisuga* and selected heteropterans were processed with OrthoFinder, and single-copy orthologs were used for phylogenetic reconstruction and divergence-time estimation. Orthogroup count matrices were used for CAFE5 gene-family expansion/contraction analyses. Single-copy orthogroups were further screened for positive selection using codon alignments and HyPhy-aBSREL and MEME models.

**S2. Mitochondrial and bacterial sequence identification**

Bacterial sequences were identified after genome assembly with BlobTools. For each genome assembly, contigs were searched against the NCBI BLAST using the following command.

blastn -query rubida.genome.fasta -db nt -out rubida.blast.txt -outfmt @ ”6 qseqid sseqid length qstart qend sstart send pident evalue” -max_target_seqs 1 -num_threads 80

blastn -query rhodnius.genome.fasta -db nt -out rhodnius.blast.txt -outfmt @ ”6 qseqid sseqid length qstart qend sstart send pident evalue” -max_target_seqs 1 -num_threads 80

Corresponding raw reads were mapped back to each assembly to generate a sorted BAM file coverage estimation. BlobTools was then used create a blobDB from the assembly FASTA file, BLAST results, and the BAM file using the following command.

minimap2 -ax map-hifi -t 80 rhodnius.genome.fasta rhodnius.hifi_reads.fq.gz | samtools sort -@ 80 -o rhodnius.sorted.bam

minimap2 -ax map-hifi -t 80 rubida.genome.fasta rubida.hifi_reads.fq.gz | samtools sort -@ 80 -o rubida.sorted.bam

blobtools create -i rubida.genome.fasta -b rubida.sorted.bam -t rubida.blast.txt -o rubida_blob

blobtools view -i rubida_blob.blobDB.json -o rubida_blob_view

blobtools create -i rhodnius.genome.fasta -b rhodnius.sorted.bam -t rhodnius.blast.txt -o rhodnius_blob

blobtools view -i rhodnius_blob.blobDB.json -o rhodnius_blob_view

Mitochondrial contigs were identified annotated using MitoHiFi and MitoFinder. Mitochondrial genomes from *T. rubida* and *R. prolixus* were used as reference input and to dtect and annotate mitochondrial sequences from both the genome assemblies using the following command.

mitohifi.py -c rubida.genome.fasta -f rubida.MT556664.1.fasta -g rubida.MT556664.1.gb -t 80 -o 5

mitohifi.py -c rhodnius.genome.fasta -f rhodnius.NC_050328.1.fasta -g rubida. NC_050328.1.gb -t 80 -o 5

***Wolbachia* genes in *R. prolixus genome***

The *R. prolixus* genome was screened for a set of 30 *Wolbachia* genes previously identified as horizontally transferred using Blastn algorithm . Additionally, for a thorough identification of any *R. prolixus* genomic regions with a putative bacterial origin, all the scaffolds from filtered assembly were annotated utilizing a prokaryote specific tool Prokka v1.14.6 . BlastN and BlastP searches for all Prokka identified features were run against nt and nr databases. The features returning any *Wolbachia* sp. as the best blast hit were considered as of putative *Wolbachia* origin. *Wolbachia* origin was verified with back BlastX searches against nr and the genes were subjected to manual curation using SnapGene® v7.1.1 software (www.snapgene.com). Pseudogenes were identified by Pseudofinder v1.1.0 . All *Wolbachia* genes were translated into protein sequences using gene2protein.py script (<https://github.com/hassantarabai/MS-Rhodnius>), then subjected to functional annotation and ortholog assignment, identifying COGs and their corresponding categories using eggNOG-mapper v2.1.12 .

Codon usage across different *Wolbachia* regions within the generated *R. prolixus* genome was analyzed and compared against the rest of the *R. prolixus* genome and different *Wolbachia* backgrounds. Three *Wolbachia* codon usage backgrounds were used: the originally calculated *Wolbachia* background used by Mesquita and colleagues, *Wolbachia* WCle, endosymbiont of *Cimex lectularius* (GenBank accession no. GCA_000829315.1), and *Wolbachia* background calculated as an average of codon usage across two newly assembled *Wolbachia* genomes associated with *R. pictipes* (details below), using EMBOSS v5.0.0 cusp module. Additionally, the comparative codon usage analysis also contained the data for originally identified *Wolbachia* inserts as described by Mesquita and colleagues.

In order to calculate *Wolbachia* codon usage background related to *Rhodnius* species, we have utilized SRA data available for *Rhodnius pictipes* (SRR6749974 and SRR6749976) for which the association with *Wolbachia* symbionts have been recently shown. The data were assembled using metaSPAdes v3.15.3 software (Nurk et al.,2017) in the Kbase environment. To identify *Wolbachia* contigs, the meta-assemblies were screened by BLASTn search (default E-value 10.0 and one best hit) against the nt database. The contigs within the final *Wolbachia* metagenome assembled genomes (MAGs) were refined based on 3 criteria: (1) BLASTn hit with 80% or higher identity to any *Wolbachia* spp., (2) hit length greater than 100 base pairs, and (3) similar coverage of the contigs passing the two previous criteria. The MAGs underwent again a BLASTn (default E-value 10.0 and one best hit) search against the nt database to validate the accurate assignment to *Wolbachia*.

Gene expression analysis of *Wolbachia* genes in *R. prolixus* used a similar methodological approach to that performed on the complete transcripts of *R. prolixus*. In addition, a heatmap was generated to visualize the individual *Wolbachia* genes expression among compared groups. This resulted in the generation a Euclidean clustering heatmap to infer the expression patterns of the integrated *Wolbachia* genes among compared datasets. (Refer to DGE_analysis.R script at <https://github.com/hassantarabai/MS-Rhodnius> for the complete R code).

For building *Wolbachia* insert phylogeny, putative *Wolbachia* insert sequences were extracted from the genome assembly and analyzed together with representative *Wolbachia* genomes using OrthoFinder to identify orthologous genes. Three single-copy orthologs were recovered and individually aligned using Clustal Omega. The alignments were concatenated, and poorly aligned regions were manually removed, resulting in a final alignment comprising 802 amino acid positions.

Phylogenetic reconstruction was performed using RAxML with the DAYHOFF amino acid substitution model and the GAMMA model of rate heterogeneity. Branch support was assessed using 100 rapid bootstrap replicates. Node labels indicate bootstrap support values, and branch lengths are proportional to the estimated number of substitutions per site.

***Arsenophonus* symbiont associated with *T. rubida***

In our *T. rubida* assembly, we identified *Arsenophonus*-like sequences. The primary purged *T. rubida* genome assembly was searched against the NCBI nucleotide database using BLASTN, and contigs with significant similarity to *Arsenophonus* were retained. A single contig of approximately 3.9 Mb was recovered, representing a near-complete *Arsenophonus* genome. The genome was annotated using PROKKA, completeness was assessed with BUSCO, and prophage regions were identified using PHASTEST.

For *Arsenophonus* phylogenetic analysis, single-copy orthologs were identified using OrthoFinder. A total of 101 single-copy orthologs, comprising 32,506 amino acid positions after concatenation, were used for phylogenetic reconstruction. Maximum-likelihood phylogenetic inference was performed using RAxML with the DAYHOFF amino acid substitution model and the GAMMA model of rate heterogeneity. Branch support was evaluated using 100 rapid bootstrap replicates. Node labels indicate bootstrap support values, and branch lengths are proportional to the estimated number of substitutions per site

**S3. Repeat discovery and genome masking**

Repeat analysis were performed for all genomes using RepeatModeler and RepeatMasker. For each genome, a species-specific RepeatModeler database was first generated using *BuildDatabase*. De novo repeat-family discovery was then performed using RepeatModeler. The de novo RepeatModeler library was combined with an invertebrate repeat library (*invrep.fasta*) derived from Repbase. The combined library was then supplied to RepeatMasker using the *-lib* function (refer to repeat analysis script at <https://github.com/Thenameisinsan/Triatominae-comparative-genomics/blob/main/repeat_modelling.py> for complete python script).

**S4. RNA-Seq filtering and genome-guided transcriptome assembly**

RNA-seq reads from different sample types of *R. prolixus* and *T. rubida* were quality filtered prior to genome alignment. Paired-end reads were processed using Trimmomatic v0.39. Overrepresented sequences identified from FASTQC reports were removed using *ILLUMINACLIP* with custom a FASTA file and reads shorter than the minimum retained length after trimming were removed with *MINLEN:36*. Only surviving paired reads were retained for downstream analyses (refer to Trimmomatic script at <https://github.com/Thenameisinsan/Triatominae-comparative-genomics/blob/main/run_trimmomatic.py> for complete script).

Filtered RNA-seq reads were aligned to their respective genome assemblies using HISAT2. RNA-seq alignments were converted to sorted BAM format using SAMtools and were used as input for genome-guided transcriptome assembly with Trinity (refer to <https://github.com/Thenameisinsan/Triatominae-comparative-genomics/blob/main/run_trinity.py> for full trinity script).

**S5. Genome annotation**

Genome annotation was performed for *R. prolixus* and *T. rubida* using NCBI’s Eukaryotic Genome Annotation Pipeline-External, EGAPx v0.4.1. For each species, a YAML configuration file was prepared containing the genome assembly, organism taxonomic identifier, and RNA-seq/transcript evidence. EGAPx was then executed through its Nextflow workflow to generate structural genome annotations. In addition, to main the annotations uniform we reannotated the *T. sanguisuga* and *N. tenuis* genome using the same EGAPx workflow and publicly available RNA-seq data from NCBI.

**YAML configurations:**

```
#nesidiocoris.yaml
genome: /home/insan/genomes/egap/nesidiocoris/input/nesidiocoris-genome.fasta
taxid: 355587
short_reads:
  - SRR5061718
  - SRR5061719
  - SRR5061720

#sanguisuga.yaml
genome: /home/insan/CompGenomics/genomes/sanguisuga.fasta
taxid: 72494
short_reads:
  - - L14l6MATES
    - - /mnt/data2/insan/virome_newData/Fasta_files/rawReads/L14l6MATES/L14l6MATES_1.fq
      - /mnt/data2/insan/virome_newData/Fasta_files/rawReads/L14l6MATES/L14l6MATES_2.fq
  - - L14l6MAGUT
    - - /mnt/data2/insan/virome_newData/Fasta_files/rawReads/L14l6MAGUT/L14l6MAGUT_1.fq
      - /mnt/data2/insan/virome_newData/Fasta_files/rawReads/L14l6MAGUT/L14l6MAGUT_2.fq
  - - L14l6FBOVARY
    - - /mnt/data2/insan/virome_newData/Fasta_files/rawReads/L14l6FBOVARY/L14l6FBOVARY_1.fq
      - /mnt/data2/insan/virome_newData/Fasta_files/rawReads/L14l6FBOVARY/L14l6FBOVARY_2.fq
  - - L14l6FBGUT
    - - /mnt/data2/insan/virome_newData/Fasta_files/rawReads/L14l6FBGUT/L14l6FBGUT_1.fq
      - /mnt/data2/insan/virome_newData/Fasta_files/rawReads/L14l6FBGUT/L14l6FBGUT_2.fq

#rubida.yaml
genome: /home/insan/genomes/egap/rubida/rubida_final_genome.fasta
taxid: 7227
short_reads:
  - - TrubFAGonads
    - - /home/insan/genomes/egap/rubida/rubida_rna/TrubFAGonads/TrubFAGonads_FKRN240281140-1A_22H3VWLT4_L7_1.fq.gz
      - /home/insan/genomes/egap/rubida/rubida_rna/TrubFAGonads/TrubFAGonads_FKRN240281140-1A_22H3VWLT4_L7_2.fq.gz
  - - TrubidaL1B
    - - /home/insan/genomes/egap/rubida/rubida_rna/TrubidaL1B/TrubidaL1B_FKRN240281131-1A_22H3VWLT4_L7_1.fq.gz
      - /home/insan/genomes/egap/rubida/rubida_rna/TrubidaL1B/TrubidaL1B_FKRN240281131-1A_22H3VWLT4_L7_2.fq.gz
  - - TrubidaL2A
    - - /home/insan/genomes/egap/rubida/rubida_rna/TrubidaL2A/TrubidaL2A_FKRN240281132-1A_22H3VWLT4_L7_1.fq.gz
      - /home/insan/genomes/egap/rubida/rubida_rna/TrubidaL2A/TrubidaL2A_FKRN240281132-1A_22H3VWLT4_L7_2.fq.gz
  - - TrubidaL3A
    - - /home/insan/genomes/egap/rubida/rubida_rna/TrubidaL3A/TrubidaL3A_FKRN240281133-1A_22H3VWLT4_L7_1.fq.gz
      - /home/insan/genomes/egap/rubida/rubida_rna/TrubidaL3A/TrubidaL3A_FKRN240281133-1A_22H3VWLT4_L7_2.fq.gz
  - - TrubidaL4A
    - - /home/insan/genomes/egap/rubida/rubida_rna/TrubidaL4A/TrubidaL4A_FKRN240281134-1A_22H3VWLT4_L7_1.fq.gz
      - /home/insan/genomes/egap/rubida/rubida_rna/TrubidaL4A/TrubidaL4A_FKRN240281134-1A_22H3VWLT4_L7_2.fq.gz
  - - TrubidaL5A
    - - /home/insan/genomes/egap/rubida/rubida_rna/TrubidaL5A/TrubidaL5A_FKRN240281135-1A_22H3VWLT4_L7_1.fq.gz
      - /home/insan/genomes/egap/rubida/rubida_rna/TrubidaL5A/TrubidaL5A_FKRN240281135-1A_22H3VWLT4_L7_2.fq.gz
  - - TrubidaMA
    - - /home/insan/genomes/egap/rubida/rubida_rna/TrubidaMA/TrubidaMA_FKRN240281142-1A_22H3VWLT4_L7_1.fq.gz
      - /home/insan/genomes/egap/rubida/rubida_rna/TrubidaMA/TrubidaMA_FKRN240281142-1A_22H3VWLT4_L7_2.fq.gz
  - - TrubidaTesM
    - - /home/insan/genomes/egap/rubida/rubida_rna/TrubidaTesM/TrubidaTesM_FKRN240281141-1A_22H3VWLT4_L7_1.fq.gz
      - /home/insan/genomes/egap/rubida/rubida_rna/TrubidaTesM/TrubidaTesM_FKRN240281141-1A_22H3VWLT4_L7_2.fq.gz
  - - TrubidDiGGMA
    - - /home/insan/genomes/egap/rubida/rubida_rna/TrubidDiGGMA/TrubidDiGGMA_FKRN240281137-1A_22H3VWLT4_L7_1.fq.gz
      - /home/insan/genomes/egap/rubida/rubida_rna/TrubidDiGGMA/TrubidDiGGMA_FKRN240281137-1A_22H3VWLT4_L7_2.fq.gz
  - - TrubidFatBL5
    - - /home/insan/genomes/egap/rubida/rubida_rna/TrubidFatBL5/TrubidFatBL5_FKRN240281136-1A_22H3VWLT4_L7_1.fq.gz
      - /home/insan/genomes/egap/rubida/rubida_rna/TrubidFatBL5/TrubidFatBL5_FKRN240281136-1A_22H3VWLT4_L7_2.fq.gz
  - - TrubidMalGL5
    - - /home/insan/genomes/egap/rubida/rubida_rna/TrubidMalGL5/TrubidMalGL5_FKRN240281138-1A_22H3VWLT4_L7_1.fq.gz
      - /home/insan/genomes/egap/rubida/rubida_rna/TrubidMalGL5/TrubidMalGL5_FKRN240281138-1A_22H3VWLT4_L7_2.fq.gz
  - - TrubidSalGL5
    - - /home/insan/genomes/egap/rubida/rubida_rna/TrubidSalGL5/TrubidSalGL5_FKRN240281139-1A_22H3VWLT4_L7_1.fq.gz
      - /home/insan/genomes/egap/rubida/rubida_rna/TrubidSalGL5/TrubidSalGL5_FKRN240281139-1A_22H3VWLT4_L7_2.fq.gz
```

For running egapx pipeline following commands were used:

```
#rubida
/home/software/egapx/ui/egapx.py -e singularity \  /home/insan/genomes/egap/rubida/rubida.yaml -o results

#nesidiocoris
/home/software/egapx/ui/egapx.py -e singularity \ /home/insan/genomes/egap/nesidiocoris/nesidiocoris.yaml -o results

#sanguisuga
/home/software/egapx/ui/egapx.py -e singularity \ /home/insan/genomes/egap/sanguisuga/sanguisuga.yaml -o results
```

**S6. Genome-collinearity analysis**

Genome collinearity among *R. prolixus*, *T. rubida*, and *T. sanguisuga* was analysed using SyMAP v5. For each species, the genome assembly FASTA file and corresponding GFF3 annotation file were imported into SyMAP through the graphical user interface on a Linux server accessed using XLaunch. Pairwise genome comparisons were performed for all three species combinations using the default SyMAP v5 alignment settings. SyMAP output files, including *blocks.txt* and *anchors.txt*, were exported and used for making pairwise and multi-species collinearity plots using custom R scripts in RStudio (refer to <https://github.com/Thenameisinsan/Triatominae-comparative-genomics/blob/main/plot_synteny.r> for synteny script).

**S7. Orthology inference and species phylogenetic reconstruction**

Protein-coding sequences from six heteropteran species along with *T. rubida* and *R. prolixus* were used for orthology analysis. Prior to OrthoFinder analysis, protein FASTA files were de-isoformed and deduplicated using a custom python script (<https://github.com/Thenameisinsan/Triatominae-comparative-genomics/blob/main/dedup_proteins.py>) and *seqkit rmdup* to select only the primary/longest isoform. Orthology was inferred using OrthoFinder v3.1.0 with DIAMOND as the sequence-search program, MAFFT as the multiple-sequence alignment tool, and IQ-TREE3 for gene-tree inference.

**OrthoFinder command used:**

/home/insan/miniforge3/envs/orthofinder/bin/orthofinder -f /home/insan/python/new/dedup_proteins/final/modified -o ortho_results/ -S diamond -M msa -A mafft -T iqtree3 -t 80 -a 56

The analysis of species-specific and unique orthogroups sets was done using a custom R script (<https://github.com/Thenameisinsan/Triatominae-comparative-genomics/blob/main/plot_upset.r>) with Gene count matrix as an input generated by OrthoFinder followed by functional annotation of orthogroups using Emapper. Among the unique orthogroups, lipocalins and perilipins were identified as relevant to triatomine biology. For the triatomine-associated lipocalin orthogroup (OG0001722), members were further examined using tissue-specific RNA-seq expression data using Salmon.

**Salmon command:**

for sample in fatBody gut ovary salglands testis; do salmon quant -i rhodnius_index/ -l A -1 /mnt/data2/insan/rhodnius_rnaSeq/${sample}_1.fq.gz -2 /mnt/data2/insan/rhodnius_rnaSeq/${sample}_2.fq.gz -p 80 --validateMappings -o quants/${sample}; done

salmon quantmerge --quants quants/* --column TPM --output rhodnius_tmp_merged. tsv

Further, additional lipocalin-like proteins from our dataset (*R. prolixus*, *T. rubida*, *T. sanguisuga*) along with those from the orthorgroup (OG0001722) expressed in salivary glands were used to build a phylogeny. In addition to lipocalin-like proteins from our dataset, additional triatomine lipocalin sequences were retrieved from NCBI for phylogenetic comparison using. MAFFT was used for multiple sequence alignment, TrimAl for trimming the alignment, and IQTREE2 was used for tree inference, using the following commands below.

mafft --auto --thread 80 lipocalin.apo.fasta > lipocalin.apo.aln.fasta

trimal -in lipocalin.apo.aln.fasta -out lipocalin.apo.trimmed.fasta -automated1

iqtree2 -s lipocalin.apo.trimmed.fasta -m MFP -B 1000 --alrt 1000 -T AUTO --prefix lipocalin.apo.trimmed

Single-copy orthologues sequences identified by OrthoFinder were used for phylogenetic tree construction. Protein sequences from each single-copy Orthogroup were aligned using MAFFT v7.525 (refer to used MAFFT script <https://github.com/Thenameisinsan/Triatominae-comparative-genomics/blob/main/run_mafft.py>). Poorly aligned regions were removed using TrimAl v1.5.rev1 with the -*automated1* option (refer to <https://github.com/Thenameisinsan/Triatominae-comparative-genomics/blob/main/run_trimal.py>). Filtered alignments were concatenated into a supermatrix using AMAS v1.0. Phylogenetic reconstruction was performed using IQ-TREE v2.2.0. Divergence-time estimation was performed using MCMCTree v4.9j with calibration points obtained from [www.timetree.org](http://www.timetree.org).

**AMAS command:**

AMAS.py convert -i concat_superMatrix.fasta -f fasta -d aa -u phylip -o concat_superMatrix_heteroptera.phy

**IQTREE2 command:**

iqtree2 -s concat_superMatrix.fasta -p partition.txt -m MFP+MERGE -bb 1000 -alrt 1000 -nt AUTO -T 80

**MCMCTREE control file:**

```
seed = 42
       seqfile = concat_supermatrix_heteroptera.phy
      treefile = input.trees
      mcmcfile = mcmc.out
       outfile = out.txt

         ndata = 1
       seqtype = 2    * 2: Amino acids (Proteins)
       usedata = 2    * PHASE 1: Set to 3. PHASE 2: Set to 2.
         clock = 2    * Independent rates
       RootAge = <4.0 * Max root age 400 MYA


         model = 2    * Use empirical AA models (LG, WAG, etc.)
         alpha = 0.5  * Enables gamma rate variation
         ncatG = 4    * Standard categories for gamma

     cleandata = 0    * 0: Keep gaps; 1: Remove gaps
        burnin = 5000 * Increased for a more stable start
      sampfreq = 20   * Increased to explore more parameter space
       nsample = 20000
```

**S8. Gene-family evolution analysis**

Gene-family evolution was analysed using CAFE v5. The orthogroup gene-count matrix generated by OrthoFinder and the ultrametric species tree generated from MCMCTree were used as input. Prior to final model fitting, an error model was applied to account for non-biological variation in gene-family counts.

cafe5 -i GeneCount.txt -t species_tree_cleaned.txt -p -e -c 60 -o error_model_run

The run produced an estimated error parameter of *epsilon = 0.0242007*. The final likelihood for the error-estimation run was *-lnL = 49733.5*, with an estimated global lambda of *0.061767237176614*. The maximum possible lambda for the topology was 0.357126, and 123 values were attempted with no rejected values. The resulting error model file, *Base_error_model.txt*, was supplied to the downstream CAFE analyses.

Two models were compared, a base model using a single global birth-death rate parameter lambda (λ) across all branches

cafe5 -i GeneCount.txt -t species_tree_cleaned.txt -p -eBase_error_model.txt -c 80 -o base_model_results

The base model produced a final likelihood of *-lnL = 50020.7*, with an estimated *λ = 0.061615032349192* and the same error parameter, *epsilon = 0.0242007*. The maximum possible lambda for the topology was 0.357126, and 56 values were attempted with no rejected values.

A gamma-rate model was then run using the same gene-count matrix, dated species tree, and error model, but allowing among-family rate heterogeneity using three discrete gamma rate categories (*-k 3*).

cafe5 -i GeneCount.txt -t species_tree_cleaned.txt -p -eBase_error_model.txt -k 3 -c 80 -o final_gamma_results

The gamma model produced a lower final negative log-likelihood than the base model *(-lnL = 45567.9* compared with -*lnL = 50020.7),* indicating a better fit to the gene-family count data. The gamma model estimated *lambda = 0.15490995036436*, *epsilon = 0.0242007*, and *alpha = 0.577523*, with a maximum possible lambda of 0.357126. In this run, 199 values were attempted and 3% were rejected. Because the gamma model provided the better fit and allowed rate heterogeneity among gene families, the output from *final_gamma_results* was used for downstream interpretation of gene-family expansions and contractions. One orthogroup, *OG0000003*, had a failure rate greater than 20% during the gamma-model run, with 75 failures, and therefore was removed from the analysis.

Orthogroups with p < 0.01 were considered significantly expanded or contracted. Significant orthogroups were functionally annotated using InterProScan v5.76-107.0 and eggNOG-mapper v2.1.13, followed by categorizing them into chemosensory, difestion, detoxification, structural-regulatory groups. For visualisation of significant orthogroups a custom R script was used. Refer to githhub for eggNOG-mapper (<https://github.com/Thenameisinsan/Triatominae-comparative-genomics/blob/main/run_emapper.py>), interproscan (<https://github.com/Thenameisinsan/Triatominae-comparative-genomics/blob/main/run_interproscan.py>), and R scripts (<https://github.com/Thenameisinsan/Triatominae-comparative-genomics/blob/main/plot_heatmap.r>).

Several evolved orthogroups identified as cytochrome P450 and chemosensory receptors were further classified on domain levels. The sequences from the odorant receptor (OR): OG0000004, gustatory receptor (GR): OG0000048, and ionotropic receptor (IR): OG0000092 were compared against the *Drosophila melanogaster* proteome using BLASTp through the VectorBase/VEuPathDB web interface. The top hits with both sequence similarity and annotation as OR/GR/IR or OR/GR/IR-like proteins were selected further for their receptors and to which ligands they respond to against the Database of Odorant Responses, DoOR 2.0.

Because insect odorant receptors evolve rapidly and ligand specificity cannot be assumed from sequence similarity alone, DoOR annotations were not interpreted as experimentally validated responses for the heteropteran receptors. Instead, the DoOR annotations were used as additional comparative functional annotation to identify whether these chemosensory receptors copies are like broad classes of volatile compounds. To support this, relevant literatures that studied some of these volatile compounds in insects were referenced in the manuscript.

Cytochrome P450-containing orthogroups were classified using HMMER against clan and family-level P450 profiles built from the Nelson Laboratory *Anopheles gambiae* P450 database using a custom bash script (refer to <https://github.com/Thenameisinsan/Triatominae-comparative-genomics/blob/main/build_cp450_hmmer.sh>).

**S9. Positive selection analysis**

Single copy orthogroups from OrthoFinder were used for positive selection analysis. For each orthogroup, protein sequences were aligned using MAFFT v7.525. Corresponding coding sequences were then converted into codon alignments using PAL2NAL v14 to preserve reading frame information. Refer to <https://github.com/Thenameisinsan/Triatominae-comparative-genomics/blob/main/run_pal2nal.py> for pal2nal script

Positive-selection analyses were performed using HyPhy v2.5.93(MP). The adaptive Branch-Site Random Effects Likelihood model, aBSREL, was used to test for episodic positive selection on individual branches of the phylogeny. aBSREL analyses were run across all branches without assigning foreground branches. P-values were corrected for multiple testing using the Holm-Bonferroni method, and corrected p < 0.05 was considered significant. In addition, MEME was used to detect site-level episodic diversifying selection across the phylogeny. Refer to <https://github.com/Thenameisinsan/Triatominae-comparative-genomics/blob/main/run_hyphy_absrel> and <https://github.com/Thenameisinsan/Triatominae-comparative-genomics/blob/main/run_hyphy_meme.py> for HyPhy-aBSREL and -MEME script, respectively.

Functional annotation for the positively selected orthogroups were based on Interproscan and Emapper analysis followed by categorization of the orthogroups based on NCBI’s cluster of orthogroups database. The *dNdS* plot was made with RStudio using the following R script <https://github.com/Thenameisinsan/Triatominae-comparative-genomics/blob/main/plot_dnds.r>.
