## Supplementary Figure S1 for "Highly contiguous genomes of *Rhodnius prolixus* and *Triatoma rubida* reveal the molecular basis of haematophagy evolution in Triatominae"

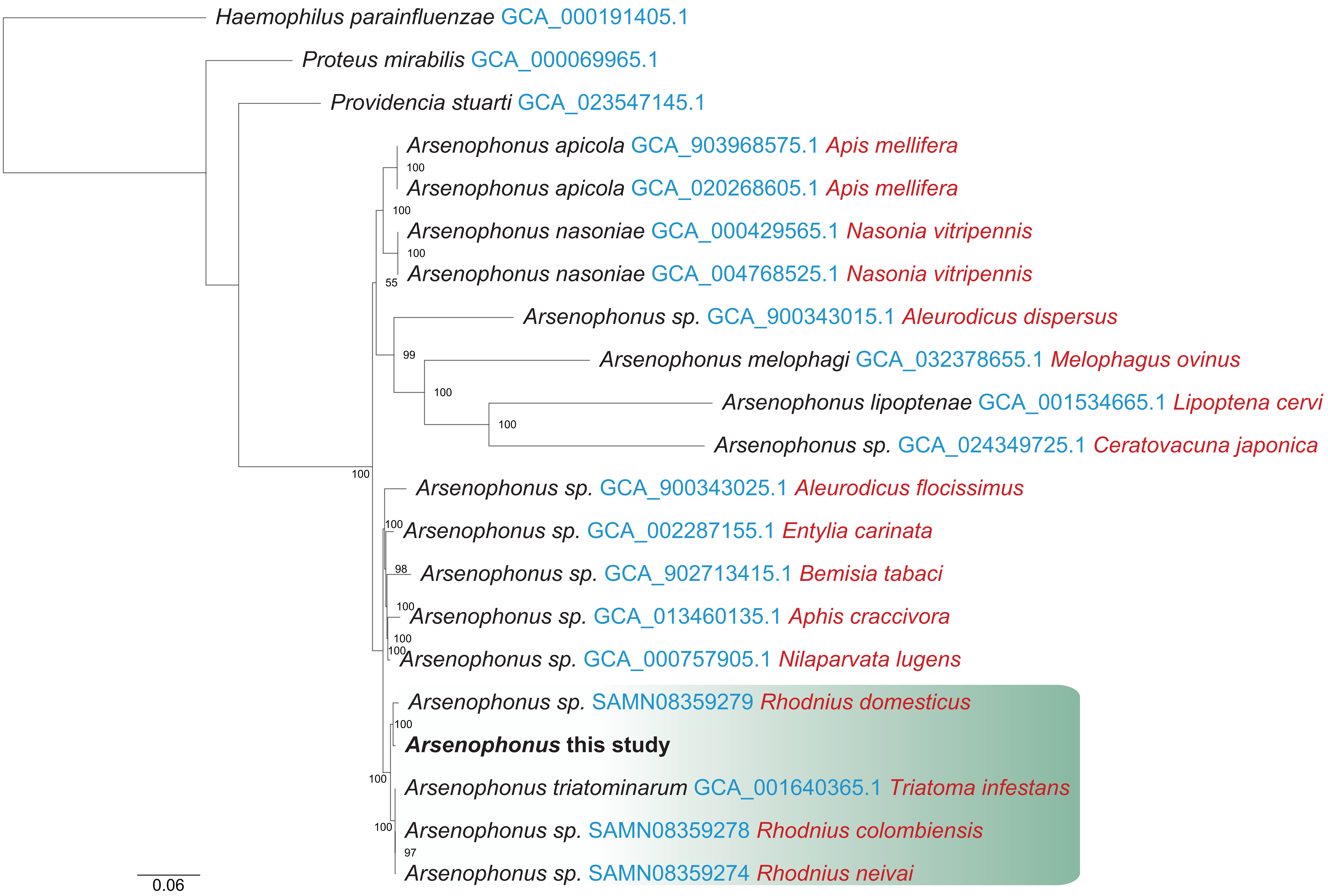


**Supplementary Figure S1. Phylogenetic placement of the *Arsenophonus* symbiont of *T. rubida*** Maximum-likelihood phylogeny showing the placement of the *Arsenophonus* sequences identified in this study relative to representative *Arsenophonus* lineages. Node labels indicate bootstrap support values above 50, and the scale bar represents substitutions per site.
