## Supplementary Figure S2 for "Highly contiguous genomes of *Rhodnius prolixus* and *Triatoma rubida* reveal the molecular basis of haematophagy evolution in Triatominae"

| Species | Genome Size | GC Content% | No. of Protein-coding Genes |
|---|---|---|---|
| *Acyrthosiphon pisum* | 533.6 Mb | 29.5 | 17,681 |
| *Halyomorpha halys* | 1.2 Gb | 31 | 14,486 |
| *Cimex lectularius* | 510.8 Mb | 35 | 11,949 |
| *Nesidicoris tenuis* | 265.6 Mb | 40 | 10,955 |
| *Rhynocoris fuscipes* | 621.6 Mb | 31.5 | 13,362 |
| *Rhodnius prolixus* | 588.46 Mb | 33.91 | 12,270 |
| *Triatoma rubida* | 934.08 Mb | 33.44 | 10,402 |
| *Triatoma sanguisuga* | 1.2 Gb | 33.5 | 11,880 |

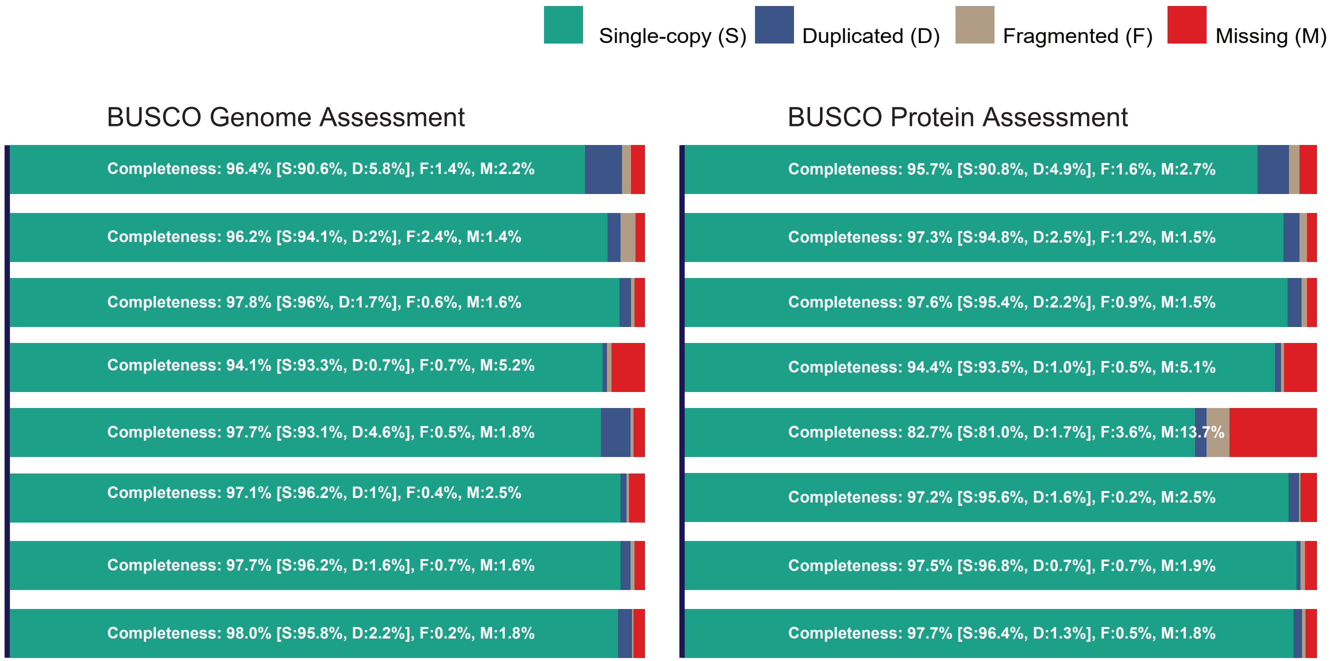


**Supplementary Figure S2. Genome assembly metrics and BUSCO genome and protein completeness assessments across analyzed insect species.** The table presents genome stats, including Genome Size, GC Content, and the Number of Protein-coding Genes for the analyzed species in this study. The completeness of genome and proteins were assessed using BUSCO v6.0.0 against the *arthropoda_odb12* lineage dataset (n=1667). The stacked bar chart shows the proportions of single copy (teal), duplicated (navy), fragmented (tan), and missing (dark red) orthologs.
