## Supplementary Figure S3 for "Highly contiguous genomes of *Rhodnius prolixus* and *Triatoma rubida* reveal the molecular basis of haematophagy evolution in Triatominae"

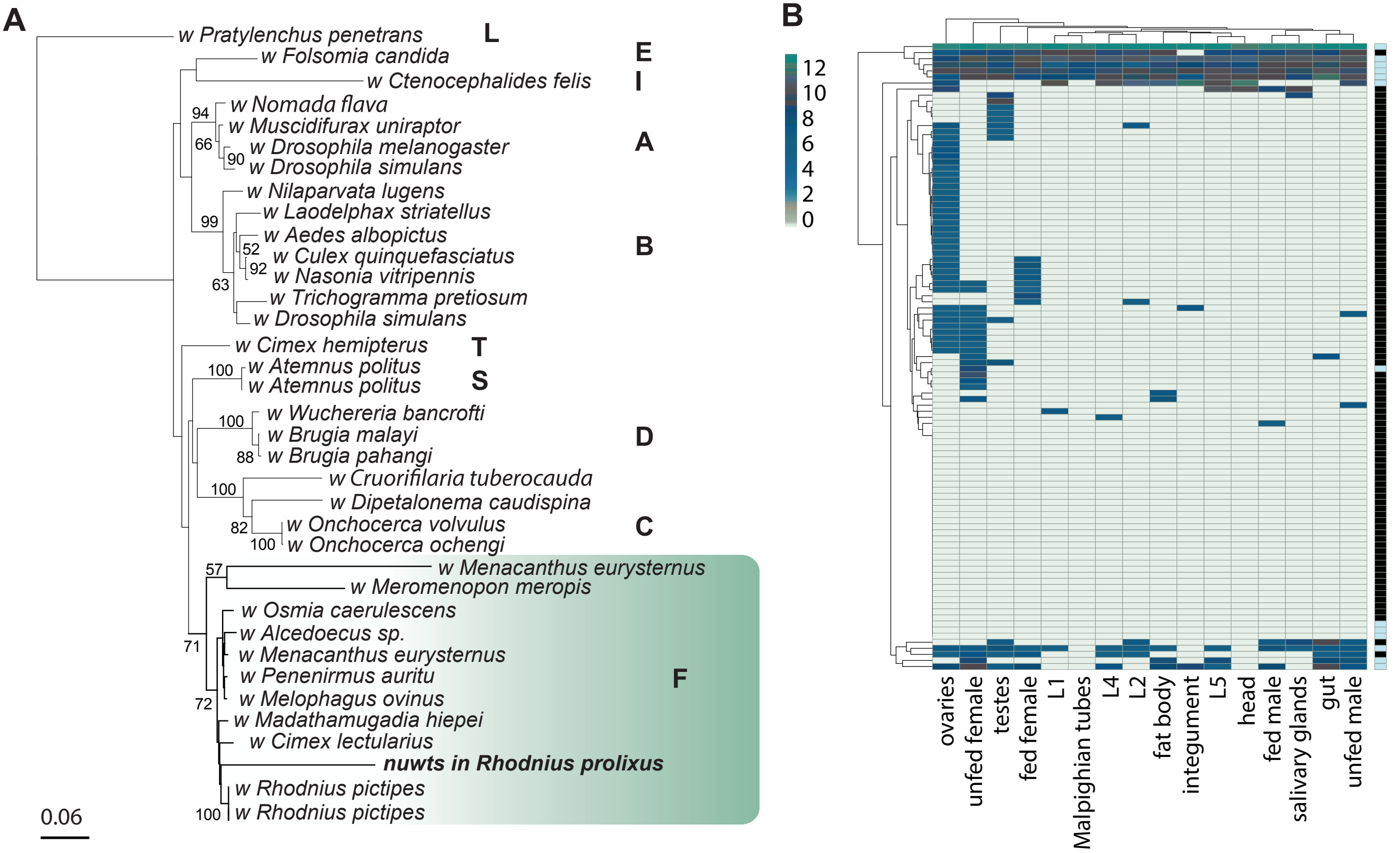


**Supplementary Figure S3. Phylogenetic placement and transcriptomic analyses of *Wolbachia*-derived genes integrated in the *Rhodnius prolixus* genome. (A)** Phylogenetic placement of the *Wolbachia*-derived insertion identified in the *R. prolixus* genome. The tree includes representative *Wolbachia* lineages from different supergroups, with the *R. prolixus Wolbachia*-derived sequence clustering within the F supergroup together with *Wolbachia* associated with other *Rhodnius* species. Node values indicate branch support, and the scale bar represents amino-acid substitutions per site. **(B)** Heatmap showing expression of integrated *Wolbachia*-derived genes across *R. prolixus* tissues and developmental stages. Rows represent expressed integrated genes, including both intact genes and putative pseudogenes, and columns represent RNA-seq samples or tissue/stage categories. Colour intensity indicates relative transcript abundance, with darker colours representing higher expression. Expression was detected for 67 of the 129 identified integrated genes, including 29 genes classified as putative pseudogenes, with the strongest overall expression observed in ovary samples.
