## Supplementary Figure S4 for "Highly contiguous genomes of *Rhodnius prolixus* and *Triatoma rubida* reveal the molecular basis of haematophagy evolution in Triatominae"

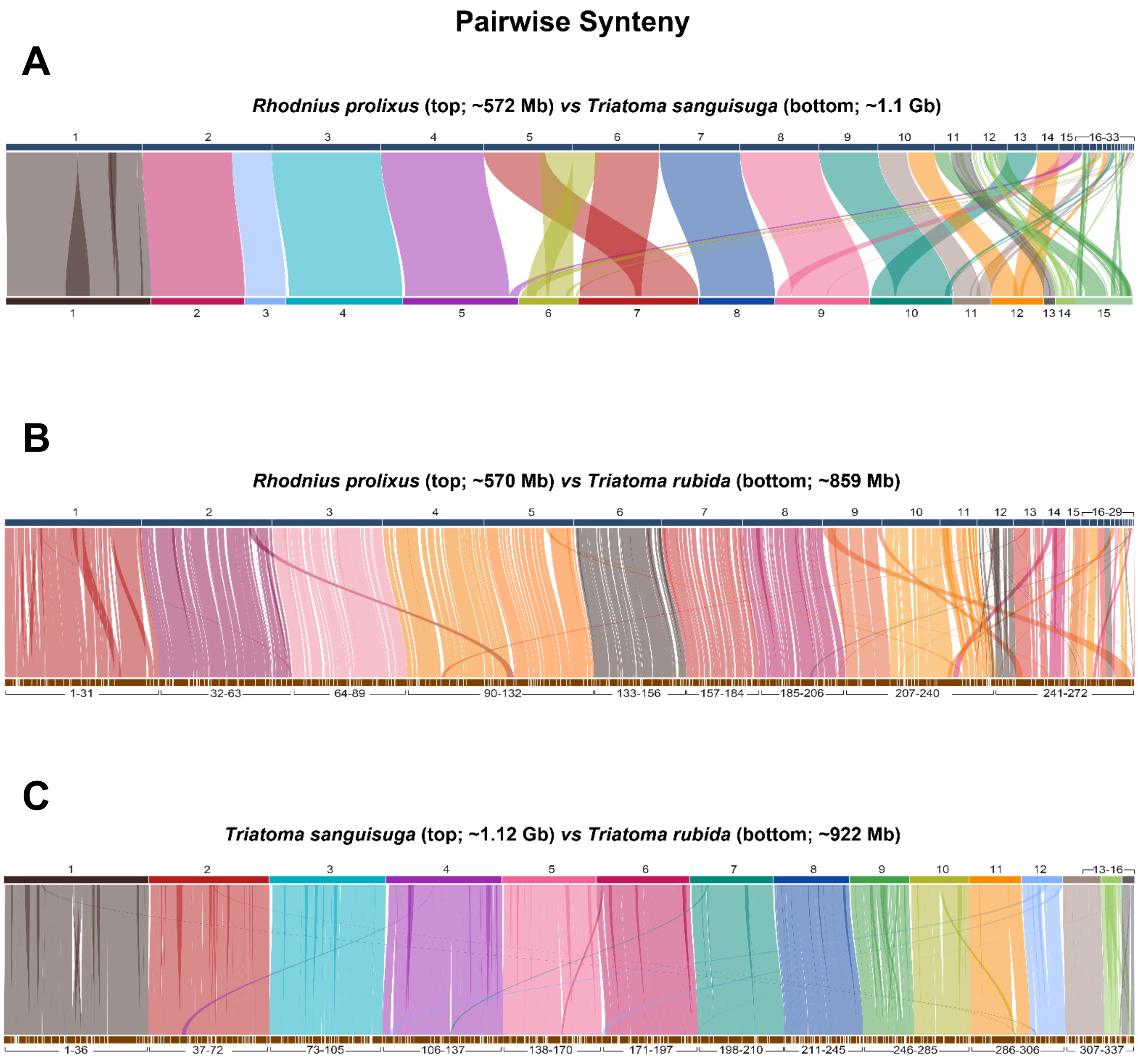


**Supplementary Figure S4. Pairwise synteny between triatomine species.** Pairwise collinearity among three triatome species: (A) *Rhodnius prolixus* and *Triatoma sanguisuga*, (B) *Rhodnius prolixus* and *Triatoma rubida*, and (C) *Triatoma sanguisuga* and *Triatoma rubida*. In each panel, species are represented as horizontal bars, with each segment corresponding to a numbered contig or scaffold (**see supplementary table**) ordered by length relative to the reference species (top bar). Coloured ribbons connect collinear blocks between species, with colours corresponding to the species with fewer contigs/scaffolds. Scaffolds included in the analysis represented 97.5% (573.8 Mb of 588.5 Mb) of the *R. prolixus* genome, 95.2% (1.11 Gb of 1.16 Gb) of the *T. sanguisuga* genome, and 98.7% (921.5 Mb of 934.1 Mb) of the *T. rubida* genome.
