## Supplementary Figure S5 for "Highly contiguous genomes of *Rhodnius prolixus* and *Triatoma rubida* reveal the molecular basis of haematophagy evolution in Triatominae"

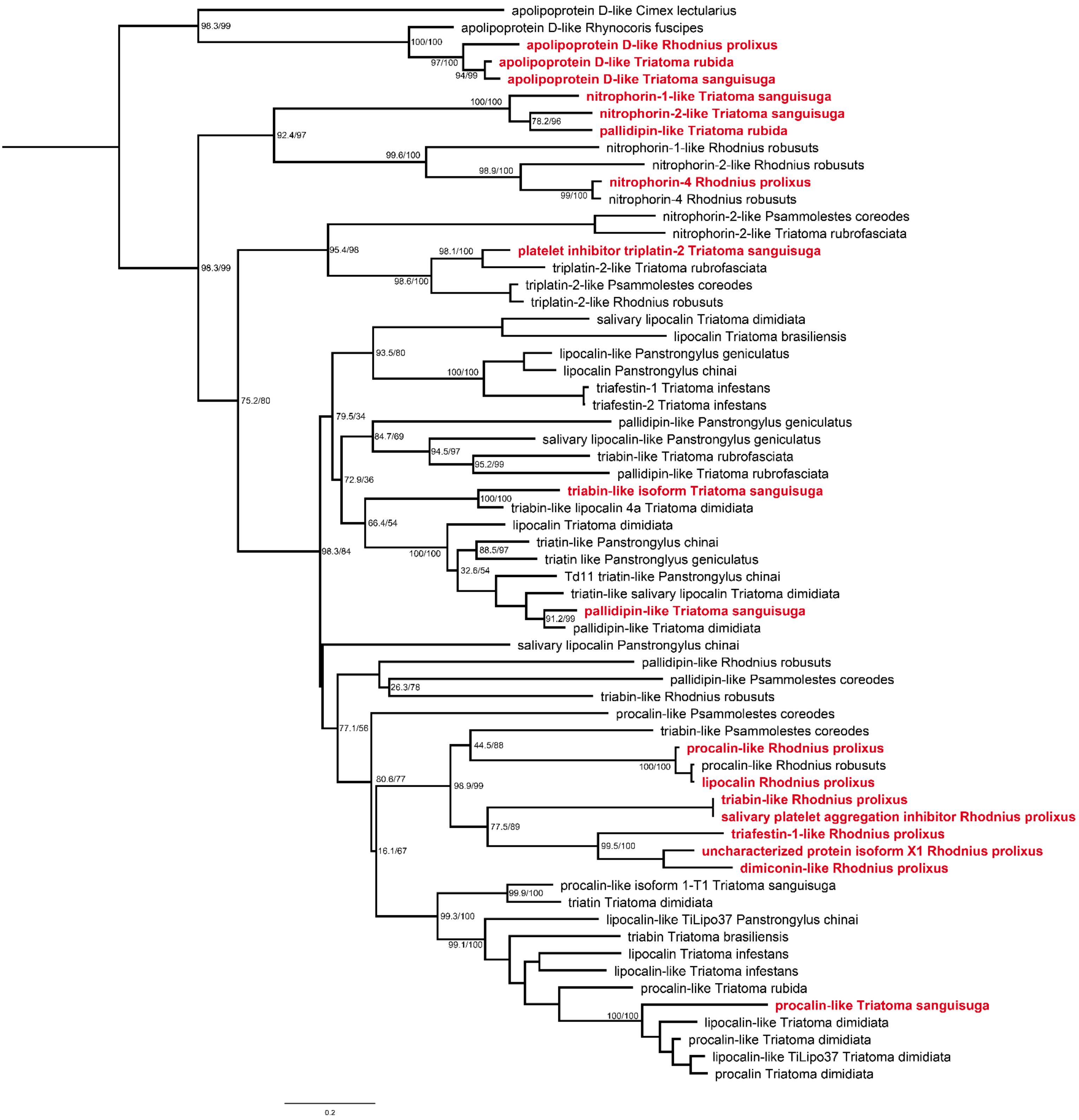


**Supplementary Figure S5. Phylogenetic placement of triatomine lipocalin-like proteins.** Maximum- likelihood tree of salivary lipocalin-like proteins identified from *Rhodnius prolixus*, *Triatoma rubida*, and *Triatoma sanguisuga*, together with representative triatomine lipocalins retrieved from NCBI. Lipocalin proteins retrieved from this study are highlighted in bold red and support values evaluated with 1, 000 ultrafast bootstrap replicated and 1, 000 SH-aLRT tests are shown under nodes.
